# Identification and characterization of new structural scaffolds modulating the activity of *Mycobacterium tuberculosis* dihydroneopterin aldolase (FolB) *in vitro*

**DOI:** 10.1101/2023.11.21.567704

**Authors:** Virginia Carla de Almeida Falcão, Alexia de Matos Czeczot, Mohammad Maqusood Alam, Kyu-Ho Paul Park, Jinyeong Heo, Minjeong Woo, Ana Micaela Camini, Luis Fernando Saraiva Macedo Timmers, David Shum, Marcia Alberton Perelló, Luiz Augusto Basso, Pablo Machado, Cristiano Valim Bizarro, Vincent Delorme

**Affiliations:** Tuberculosis Research Laboratory, Institut Pasteur Korea, Seongnam, Gyeonggi, Republic of Korea; Instituto Nacional de Ciência e Tecnologia em Tuberculose, Centro de Pesquisas em Biologia Molecular e Funcional, Pontifícia Universidade Católica do Rio Grande do Sul, Porto Alegre, Brazil; Programa de Pós-Graduação em Medicina e Ciências da Saúde, Pontifícia Universidade Católica do Rio Grande do Sul, Porto Alegre, Brazil; Chemistry Platform, Institut Pasteur Korea, Seongnam, Gyeonggi, Republic of Korea; Screening Discovery Platform, Institut Pasteur Korea, Seongnam, Gyeonggi, Republic of Korea; Programa de Pós-Graduação em Biotecnologia e Universidade do Vale do Taquari, Lajeado, Brazil; Programa de Pós-Graduação em Ciências Médicas e Universidade do Vale do Taquari, Lajeado, Brazil; Programa de Pós-Graduação em Biologia Celular e Molecular, Pontifícia Universidade Católica do Rio Grande do Sul, Porto Alegre, Brazil

**Author notes:** Neuroscience Research Institute, Gachon University, Incheon, Republic of Korea (Mohammad Maqusood Alam). R&D Center, Insol Co., Ltd., Hanam, Gyeonggi, Republic of Korea (Vincent Delorme).

**Keywords:** *Mycobacterium tuberculosis*, dihydroneopterin aldolase (DHNA), Rv3607c (FolB), Folate precursor synthesis, drug screening

## Abstract

Antifolates were among the first broad-spectrum compounds used as antimycobacterial agents and can still be of use when no other therapeutic options are available. The discovery of compounds targeting this essential pathway could lead to new therapeutic agents to treat tuberculosis (TB). In particular, the enzyme required for the conversion of 7,8-dihydroneopterin (DHNP) to 6-hydroxymethyl-7,8-dihydropterin (HP) and glycolaldehyde (GA) in the folate pathway (*Mtb*FolB, a dihydroneopterin aldolase - DHNA, EC 4.1.2.25), has received little attention as a potential drug target. Here, we conducted a small-scale diversity screening to identify *Mtb*FolB inhibitors using a microplate-based enzyme inhibition assay. About 6,000 compounds were assembled for the screening and 19 hits were identified, spanning 5 independent clusters. These compounds were tested in dose-response studies and active compounds selected for kinetic inhibition and time-dependent inhibition studies, leading to compounds with IC_50_ values ranging from 2.6 to 47 µM. A preliminary structure activity analysis was performed, revealing that bi-sulfonamide compounds could be explored for further optimizations. Docking studies highlighted two modes of binding for pyrazol-3-one compounds and, for the sulfonamide series, indicated several interactions with the catalytic Tyrosine-54 (Tyr54D) and Lysine-99 (Lys99A) residues of *Mtb*FolB. The sulfonamide compound **13** represents the first identified compound directed against *Mtb*FolB with an antimycobacterial activity.

## Introduction

Folates are essential cofactors participating in key biosynthetic processes in both prokaryotic and eukaryotic cells, notably the synthesis of methionine, thymine and purine bases. While a folate transport system based on membrane-associated proteins exist in mammals for the salvage of folates [1], most micro-organisms, including *Mycobacterium tuberculosis* (*Mtb*), must synthesize them *de novo*. This requirement led to the discovery of small molecules inhibiting the folate biosynthesis pathway (**Figure 1**), in particular the sulfonamides [2] and diaminopyrimidines [3], leading to an arrest of cellular growth. Such molecules, termed antifolates, were among the first broad-spectrum, synthetic antibiotics and have been widely used for the treatment of urinary tract infections [4], as well as leprosy, caused by *Mycobacterium leprae* [5]. The treatment of tuberculosis (TB) with synthetic drugs also started as early as 1946 with *para*-aminosalicylic acid (PAS) [6], a pro-drug incorporated in the folate pathway and leading to its downstream inhibition [7–10]. Despite these successes, antifolates are now largely absent from the modern treatment of TB, as more potent, bactericidal drugs are available. The emergence of drug-resistant infections however warrants their usage when no other therapeutic options are available [11].

**Figure 1.**
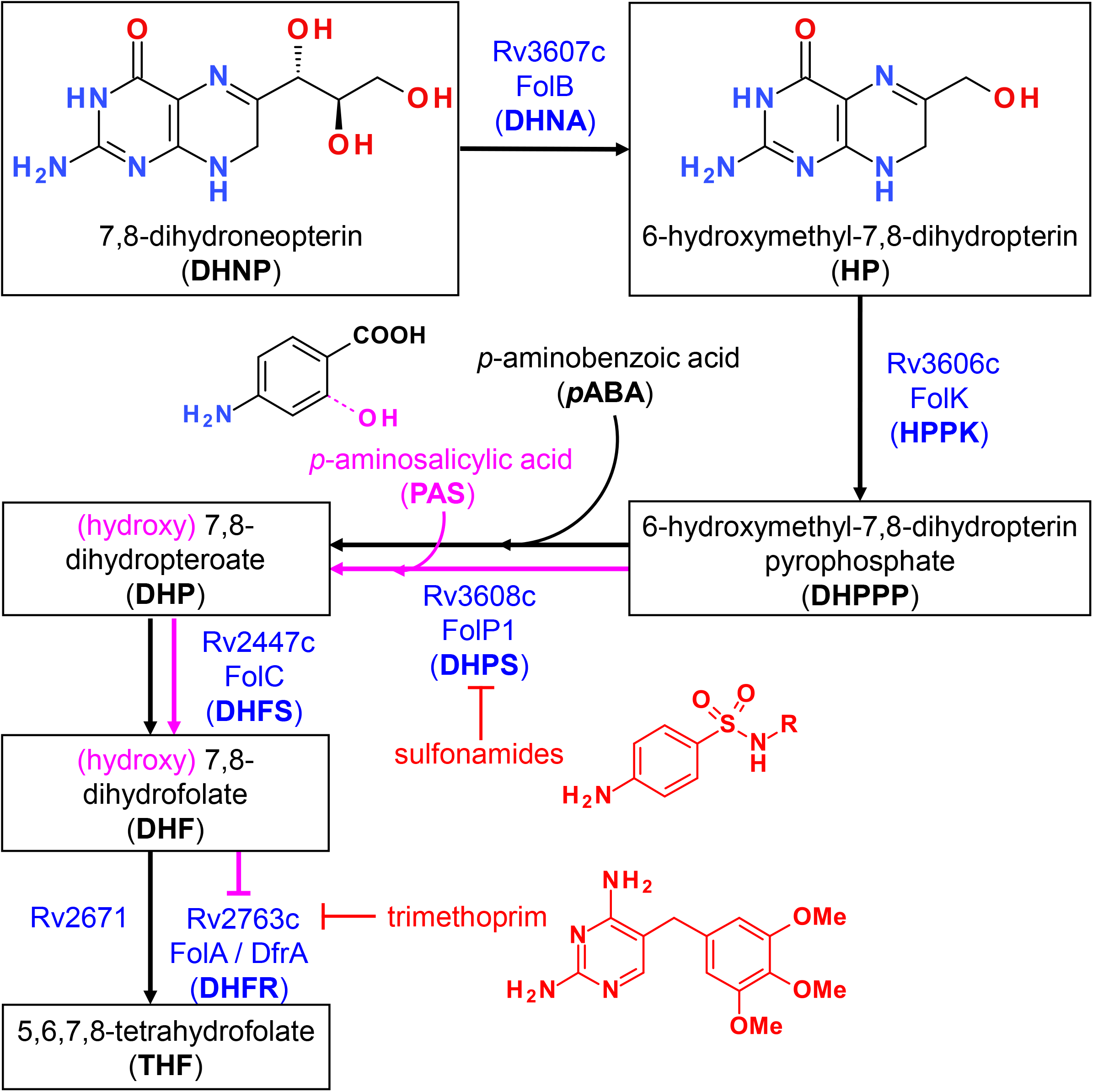
Overview of the folate synthesis pathway in *M. tuberculosis*. Steps leading to the formation of 7,8-dihydroneopterin (DHNP) are omitted for clarity. The name of the enzyme(s) proposed to catalyze each reaction is indicated in blue, along with known inhibitors in red. Incorporation of *p*-aminosalicylic acid (PAS) instead of *p*-aminobenzoic acid (PABA) at the third depicted step leads to the formation of a hydroxylated intermediate that is propagated downstream within the pathway, as indicated in pink. The metabolite ultimately inhibits FolA but not Rv2371, which has been recently identified as having a dihydrofolate reductase (DHFR) activity. Trimethoprim however is able to inhibit both enzymes at this step.

Antifolates as antibacterials have so far mostly been designed to target dihydropteroate synthase (DHPS; EC 2.5.1.15) and dihydrofolate reductase (DHFR; EC 1.5.1.3) due to their attractive active site features well conserved across bacterial species and, for DHPS, the absence of mammalian counterparts [12]. The availability of crystal structures for the two orthologues in *Mtb* (*Mtb*DHPS, Rv3608c [13]; *Mtb*DHFR, Rv2763c [14]) also greatly promoted new drug design and discovery of inhibitors with whole-cell activities against this bacteria, particularly for *Mtb*DHFR [15, 16]. The fact that Rv2671 (wrongly annotated as a riboflavin biosynthesis protein, RibD) also displays DHFR activity, although to a lower extent than *Mtb*DHFR, may however highlight additional challenges to overcome for the success of such inhibitors [17, 18]. In contrast, the catalytic reactions upstream *Mtb*DHPS assigned to Rv3606c (HPPK or FolK; EC 2.7.6.3) and Rv3607c (DHNA or FolB; EC 4.1.2.25) (**Figure 1**), have received little attention so far. In a preliminary work conducted to identify inhibitors of FolB in *Staphylococcus aureus*, guanine analogues (triazolo[4,5-d]pyrimidin-7-one) were found to display submicromolar inhibition *in vitro* despite absence of antibacterial activity [19]. Based on the results of this work and others, 8-mercaptoguanine analogues were also recently designed and tested against *Mtb*FolB, revealing also submicromolar *in vitro* activities but absence of whole cell activity [20].

Previous work highlighted distinctive features of *Mtb*FolB compared to other bacterial orthologues, in that it can catalyze an oxygenase reaction leading to the formation of 7,8-dihydroxanthopterin (DHXP) in addition to the conventional aldolase and epimerase reactions, without the need of metals or cofactor [21]. The recent confirmation that *Mtb*FolB aldolase/epimerase activity is essential for bacterial growth [22], together with this body of preliminary work, prompted us to conduct additional screening to identify inhibitors of *Mtb*FolB with greater structural diversity and less resemblance to purines. Here, we report the results of a small-scale diversity screening conducted on *Mtb*FolB and describe the initial structure-activity relationship results obtained for selected hit scaffolds.

## Materials & Methods

### Reagents and chemicals

A library of 5,674 compounds from ChemDiv (diversity set) and 400 compounds from Prestwick (F2L library) were assembled for the screening, arrayed in 96-well plates at concentration of 20 mM in DMSO and stored at -20°C. When available, the hits selected for dose-response validation were repurchased from the same suppliers. The 7,8-dihydro-D-*erythro*-neopterin (DHNP) substrate for *Mtb*FolB was purchased from Sigma-Aldrich. The 6-hydroxymethyl-7,8-dihydropterin hydrochloride (HP) substrate and the 7,8-dihydroxanthopterin (DHXP) reaction by-product were both purchased from Schircks Laboratories.

### Cloning, expression and purification of MtbFolB

The coding region of *Mtb*FolB (Rv3607c; NCBI RefSeq GeneID: 885345) was amplified by PCR using genomic DNA from *Mtb* H37Rv as a template, using primers 5’-TTTT<u>CCATGG</u>CTGACCGAATCGAA-3’ (Forward) and 5’-TTTT<u>AAGCTT</u>CATACCGCGCCGC-3’ (Reverse), and cloned in pET28a(+) vector using the NcoI/HindIII restriction sites (underlined in the primer sequences). The resulting plasmid pET28::*folB*, allowing over-expression of the native *Mtb*FolB without tags, was transformed into competent *E. coli* BL21 Star (DE3) cells (Invitrogen). Transformants were selected on LB agar plates supplemented with 50 μg/mL kanamycin. Single colonies were grown overnight at 37°C in LB medium (50 mL) supplemented with 50 μg/mL kanamycin. Part of the culture (8.5 mL) was inoculated into 500 mL of LB medium supplemented with 50 μg/mL kanamycin and grown at 37°C in shaker at 180 rpm until reaching an optical density (OD_600 nm_) of 0.4-0.6. The protein expression was induced by addition of 0.1 mM isopropyl-β-D-thiogalactoside (IPTG, Sigma-Aldrich) for 6 hours at 37°C. The cells were then harvested by centrifugation at 11,800 × *g* for 30 min at 4°C. Approximately 2 g of cell paste was resuspended in 10 mL of buffer A (25 mM Tris-HCl pH 8.0, 50 mM NaCl, 5% glycerol) containing a protease inhibitor cocktail tablet (Complete EDTA-free, Roche Diagnostics) and gently stirred for 30 min in the presence of 0.2 mg/mL lysozyme (Sigma Aldrich). Cells were subsequently disrupted by sonication (ten pulses of 10s, with cooling intervals of 1 min, at 60% amplitude) and the lysate was centrifuged at 38,900 × *g* for 30 min at 4°C. The supernatant was treated with 1% (w/v) streptomycin sulphate (final concentration) for 30 min to precipitate nucleic acids and ribonuclear proteins [23] under slow agitation and centrifuged at 38,900 × *g* for 30 min at 4°C. After centrifugation, the supernatant was dialyzed against buffer A, and loaded onto a DEAE Sepharose CL6B anionic exchange column (GE Healthcare) equilibrated with buffer A. The column was washed with 3 column volumes (CV) and adsorbed protein were eluted by a linear gradient (0-1 M NaCl) with 14.3 CV in buffer B (25 mM Tris-HCl pH 8.0, 1M NaCl, 5% glycerol) at a flow rate of 1 mL/min. The fractions eluted from anionic exchange column were incubated with ammonium sulfate to a final concentration of 1M and loaded on a Butyl Sepharose High Performance HiLoad16/10 (GE Healthcare) equilibrated with buffer C (25 mM Tris-HCl pH 8.0, 1M ammonium sulfate, 5% glycerol). The column was washed with 7 CV of buffer C and the adsorbed proteins were eluted by a linear gradient (1 to 0 M ammonium sulfate) with 20 CV in buffer C at a flow rate of 1 mL/min. The fractions containing the enzyme were pooled, concentrated using an Amicon Ultra centrifugal filter with a 10 kDa cut-off (Millipore) and loaded on a HiLoad Superdex 200 26/60 size exclusion column (GE Healthcare) previously equilibrated with buffer A. Protein was isocratically eluted with 1 CV of buffer A at a flow rate of 0.3 mL/min. Fractions containing pure FolB were pooled and concentrated to 0.4-0.6 mg/mL, aliquoted and flash-frozen for conservation at -80°C.

### Microplate-based enzyme inhibition assay and screening

All enzymatic reactions were conducted in 384-well plates (Greiner, Cat #781900) using buffer A (25 mM Tris-HCl pH 8.0, 50 mM NaCl, 5% glycerol), in presence of 5 μM 7,8-dihydroneopterin (DHNP, Sigma-Aldrich) and 10 ng of recombinant *Mtb*FolB (14 nM) in a total volume of 50 μL. Incubations were done at room temperature (24°C) for 4h and the end-point fluorescence reading obtained with a multimode microplate reader (Envision or EnSight, Perkin Elmer) equipped with excitation 405 nm and emission 535 nm filters. Each assay plate included 4 full columns of controls: 2 columns for the positive control (1% DMSO) and 2 columns for the negative control (heat-inactivated *Mtb*FolB), to assess the assay performance. For the negative control, an aliquot of the pure *Mtb*FolB preparation used for the assay was taken and inactivated at 95°C for 10 min the same day.

For the screening, a fresh powder aliquot of substrate (DHNP) was added to buffer A to reach a concentration of 5 mM and solubilized by water-bath sonication for 1h in the dark. This stock was further diluted in buffer A to prepare a working solution at 12.5 μM. Stock solution of compounds in DMSO were diluted in buffer A and added to the plate using an automated dispenser (10 μL), followed by the substrate (20 μL) and the enzyme (20 μL).

Compounds were first diluted at 2 mM in 100% DMSO and 0.5 µL of this intermediate dilution was added to the assay plate, to reach a final concentration of 20 μM and a residual DMSO concentration of 1%. The fluorescence values obtained for each compound were normalized with that of the controls for the corresponding plates, using the average value for the positive control (1% DMSO) as reference for 0% inhibition and the average value of the negative control (heat-inactivated *Mtb*FolB) as reference for 100% inhibition.

### Dose-response assay (end-point assay)

For determination of compound activity by dose-response, a similar procedure was followed using compound concentration starting at 100 μM and a total of 10 doses with 2-fold dilution between each dose, with 1% DMSO final concentration in each well. Each dose-response was performed as a technical duplicate. Raw fluorescence values were plotted as a function of the compound concentration, expressed in log scale, and the data fitted against a sigmoidal dose-response model (4 parameters) using the software Prism v6 (GraphPad). The concentration required to inhibit the enzyme activity by 50% (IC_50_) was obtained as the best fit value for the IC_50_ parameter when the fitting converged and the 95% CI are reported. When necessary (absence of low plateau), constraints were added to the bottom value of the dose-response, constraining the value of this parameter above or equal to that of the negative control (inactivated enzyme). In such cases (no plateau), the 95% CI values were omitted (interval too large).

### Kinetic assay

Enzyme inhibition studies were performed using a RF-5301 spectrofluorometer (Shimadzu), monitoring an increase in fluorescence, corresponding to the formation of HP, with excitation wavelength at 365 nm and fluorescence emission at 525 nm. The slits were 10 nm for excitation and 15 nm for emission [20]. Reactions were conducted during 6 min at 25 °C using buffer A (25 mM Tris-HCl pH 8.0, 50 mM NaCl, 5% glycerol), in presence of 1.5 μM DHNP (non-saturating concentration) and 300 nM of recombinant *Mtb*FolB in a total volume of 500 μL (quartz cuvette). The DMSO concentration was fixed at 10% and the maximal rate of enzymatic reaction (100% of *Mtb*FolB activity) was determined in these conditions, in absence of inhibitor. As controls, buffer, substrate, inhibitor and enzyme spectra were performed under the same conditions to subtract fluorescence intensities that were not coming from the reaction product. Reaction velocities (μM.min^-1^) were determined from the fluorescence kinetic measurement by fitting the data points (20) to a straight line by linear regression, using compound concentrations ranging from 50 μM to 1 mM. For each compound, velocities were then plotted as a function of concentration and the plot fitted by non-linear regression using the equation below:

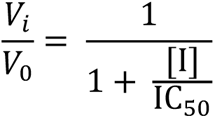

in which *V*_i_ and *V*_0_ are, respectively, the reaction velocity in the presence and in the absence of inhibitor (I).

### Time-dependent enzyme inhibition

Prior to activity determination, *Mtb*FolB (300 nM) was incubated at room temperature in buffer A together with the compound (**3**, **9** or **20**). At the corresponding time-point (0, 30 and 60 min), the substrate was added, and the enzyme velocity determined as described above (kinetic assay). Different incubation and test conditions were used. *i*) Normal condition: *Mtb*FolB was incubated with the compound at 300 μM and 2% DMSO, then the velocity measured in presence of the compound (300 μM), 1.5 μM of substrate and 2% DMSO. *ii*) Excess of substrate: *Mtb*FolB was incubated with the compound at 300 μM and 2% DMSO, then the velocity measured in presence of the compound (300 μM), 3 μM (2×) of substrate and 2% DMSO. *iii*) Excess of inhibitor: *Mtb*FolB was incubated with the compound at 3 mM (10×) and 15% DMSO, then the velocity measured in presence of the compound (300 μM) and 1.5 μM of substrate with DMSO at a final concentration of 2%.

### Chemistry

All procedures for the synthesis of the hit of interests and their derivatives, along with NMR and mass spectrometry characterization data, are available in the supplementary methods of this article.

### Docking studies

All docking procedures were performed using AutoDock4.2 program. To ensure that all compounds were properly docked, a 3D-grid with dimensions 50 × 50 × 50 with spacing of 0.375 Å was used to define the active site. The grid was defined in the 6-hydroxymethyl-7,8-dihydropterin (HP) binding region, which is located at the interface of two subunits, based on the octameric structure of *Mtb*FolB obtained by crystallography (PDB: 1NBU) [24]. A re-docking procedure was performed to evaluate whether the program could reproduce the ligand location found in the structure. The Lamarckian Genetic search Algorithm was used with 60 runs, and the other parameters were set to their default values, except for the *number of evaluations* and the *number of individuals in population*, which were set to 4,000,000 and 450, respectively. Docking results shown in **Figures 4 and 5** were prepared using PyMOL v2.5.3 (DeLano Scientific).

### Mycobacterium tuberculosis inhibition assay

The activity of the compounds **3**, **9**, **13**, **16**, **20**, **21**, **22**, and **23** was evaluated against *M. tuberculosis* H37Rv reference strains (ATCC 27294) using a colorimetric assay to determine their minimum inhibitory concentration (MIC), as previously described [25]. Isoniazid (MIC = 2.3 µM) and rifampicin (MIC = 0.2 µM) were used as controls in this experiment. The MIC determination procedures were carried out in Difco^TM^ Middlebrook 7H9 broth (Becton Dickinson – BD) supplemented with 10% (v/v) BBL^TM^ Middlebrook ADC enrichment (albumin, dextrose and catalase – BD) at a final concentration of DMSO equal to 2.5%. The compounds were evaluated using 8-point, 2-fold serial dilutions, with the maximum concentration being 200 µM for the compounds **3**, **9**, **16**, **20**, **21**, **22**, **23** and 20 µg/mL (equivalent to 55.4 µM) for the compound **13**. Three independent experiments were conducted for each compound, and the MIC was considered as the most frequent value among the three obtained.

## Results and Discussion

### Assay development, screening and hit confirmation

Based on previous work performed using FolB from *S. aureus* [19] and *Mtb* [20, 21], we established an *in vitro* enzymatic assay for the evaluation of compound libraries arrayed in 384-well plates. The assay is based on the fluorescence of 6-hydroxymethyl-7,8-dihydropterin (HP), the kinetic product formed by action of *Mtb*FolB on the substrate 7,8-dihydroneopterin (DHNP). The fluorescence of the mixture containing the substrate alone was found to be stable, even after 20h incubation (**Figure S1A**). The thermodynamic product, 7,8-dihydroxanthopterin (DHXP), has a weaker fluorescence intensity at the wavelength used (**Figure S1B**) and is formed at a smaller rate [21], allowing for a stable end-point measurement in a time-window from 3h and up to 20h (**Figure S1C**). The 4h time-point was selected for the screening to ensure maximum conversion of the substrate while minimizing formation of DHXP.

The assay developed was then used to screen a small library of diverse molecules (total 6,074 compounds) and the fluorescence results normalized to that of the controls on a plate-by-plate basis (**Figure 2**). Hits were selected based on a 30% inhibition threshold, yielding 19 compounds that were selected for dose-response studies (**Figure S2**). These molecules could be grouped into 5 independent clusters (numbered I-V), except for 4 compounds that were singletons. Hits were re-purchased or re-synthesized and tested at 10 different doses in duplicate using the same assay, with concentrations starting at 100 μM. Results are summarized in **Table 1**. Based on these confirmation data, clusters I, IV and V were eliminated due to lack of activity. The activity of the singletons **2**, **14** and **18** were confirmed, but we could not re-test compound **17** (not available to re-purchase). Clusters II and III were also found to be active and we decided to prioritize our efforts on the characterization of these two clusters.

**Figure 2.**
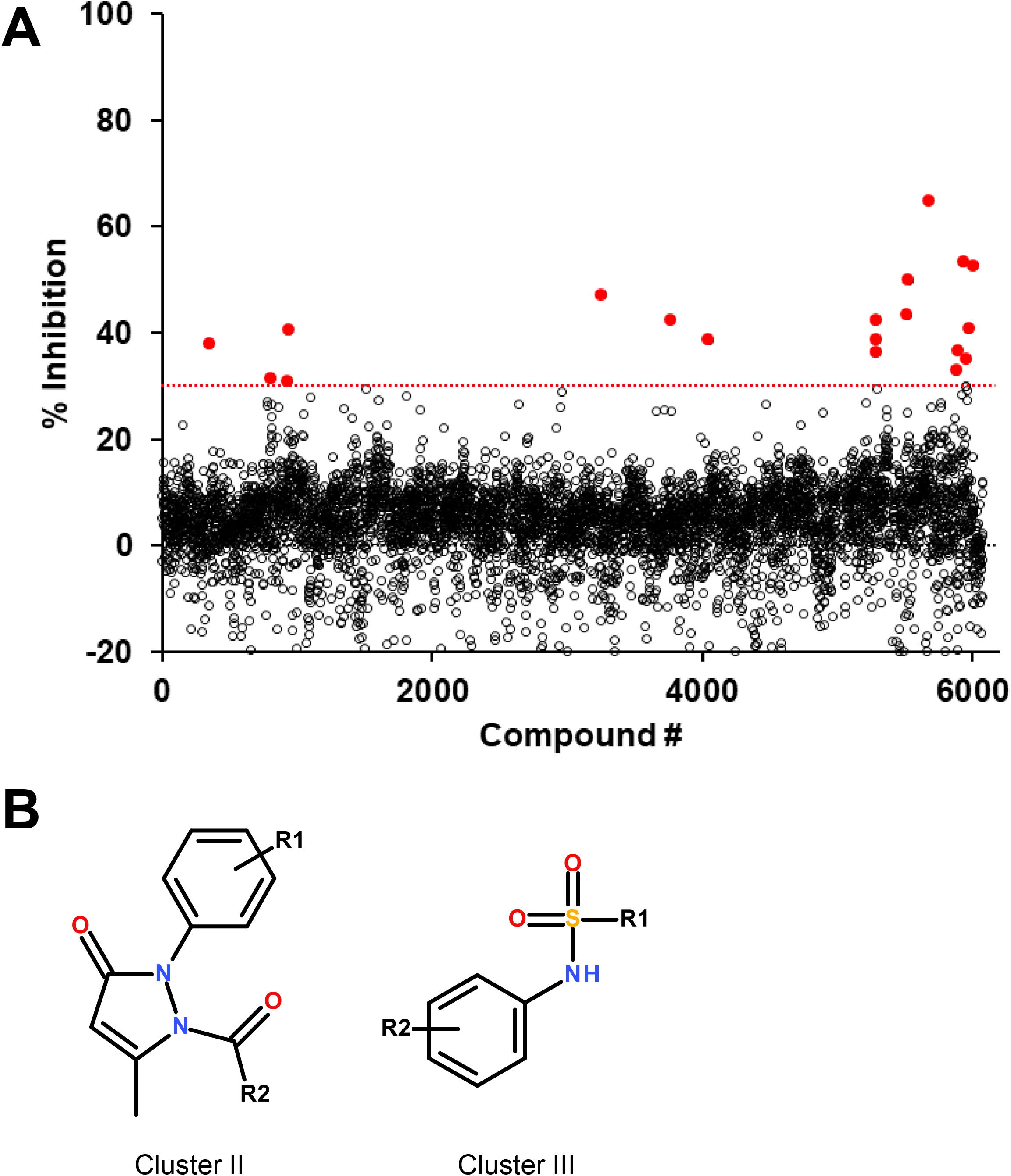
Screening results and selected hit scaffolds. **A**. Summary of the percent inhibition obtained during the primary screening, for each compound tested. The threshold for hit selection was fixed at 30% and is indicated by the dotted red line. Compounds above the threshold are indicated in red. **B**. Core structure of the two hit scaffolds that were retained following validation by dose-response.

**Table 1.** Summary of the hits selected at 30% inhibition threshold and their validation by dose-response.

| # | % inhibition | IC<sub>50</sub> (μM)<sup>a</sup> | Purchase (P) / Synthesis (Sy) | Cluster<sup>b</sup> |
| --- | --- | --- | --- | --- |
| 1 | 64.9 | NA | Sy | I |
| 2 | 53.2 | 8.0 (4.0 – 16.1)<br>Plateau inhibition (20-30%) | P | S |
| 3 | 52.5 | 30.1 (9.9 – 92.1) | P | II |
| 4 | 49.8 | NA | Sy | I |
| 5 | 47.0 | 2.6 (2.3 – 2.9) | P | III |
| 6 | 43.3 | NA | Sy | I |
| 7 | 42.4 | 63.2 (no plateau) | P | III |
| 8 | 42.2 | NA | Sy | IV |
| 9 | 40.8 | 46.8 (no plateau) | P | II |
| 10 | 40.5 | NA | Sy | V |
| 11 | 38.7 | 79.6 (no plateau) | P | III |
| 12 | 38.6 | NA | Sy | IV |
| 13 | 37.9 | 15.4 (12.0 – 19.8) | Sy | III |
| 14 | 36.6 | 17.2 (3.3 – 89.1)<br>Plateau inhibition (20-30%) | P | S |
| 15 | 36.3 | NA | Sy | IV |
| 16 | 35.1 | 17.6 (6.2 – 50.5) | P | II |
| 17 | 32.9 | NT | - | S |
| 18 | 31.5 | 13.7 (7.1 – 26.4)<br>Plateau inhibition (30-40%) | P | S |
| 19 | 31.0 | NA | Sy | V |

a. Concentration required to inhibit 50% of FolB activity. Values are best-fit for a duplicated dose-response. The 95% CI is indicated in parenthesis, except for cases where it was very wide (no plateau for the bottom value of the curve). NA: not active or IC<sub>50</sub> > 100 μM. NT: not tested.
b. Clusters were numbered from I to V, in order of appearance. S: Singleton. All structures are depicted on **Figure S2**.

### Pyrazol-3-one series (Cluster II)

The 3 active hits for this series (**3**, **9** and **16**) had IC_50_ values ranging from 18 to 47 μM in the dose-response assay. Interestingly, 4 other compounds from the initial screening library displayed similar structures and, although not selected from the primary screening, had percentages of inhibition close to the 30% threshold. These 4 compounds were re-purchased and tested in dose-response, revealing that all were active with IC_50_ values within the same range as the 3 selected hits (**Table 2**). Kinetic inhibition studies also indicated a dose-dependent inhibition of the reaction rate (**Figure S3**), but higher doses of compounds were required to see a reduction in velocity during the first minutes of the reaction, as shown by the higher IC_50_ values obtained (**Table 2**). Time-dependent inhibition studies conducted with compounds **3**, **9** and **20** indicated that the inhibition of the reaction rate was stable in time for at least 1h, reaching 40 to 60% inhibition at 300 μM (**Figure S4**). Pre-incubating the enzyme with higher doses of compounds (up to 3 mM) did not affect the inhibition of the reaction, nor did higher substrate concentrations (**Figure S4**).

**Table 2.**
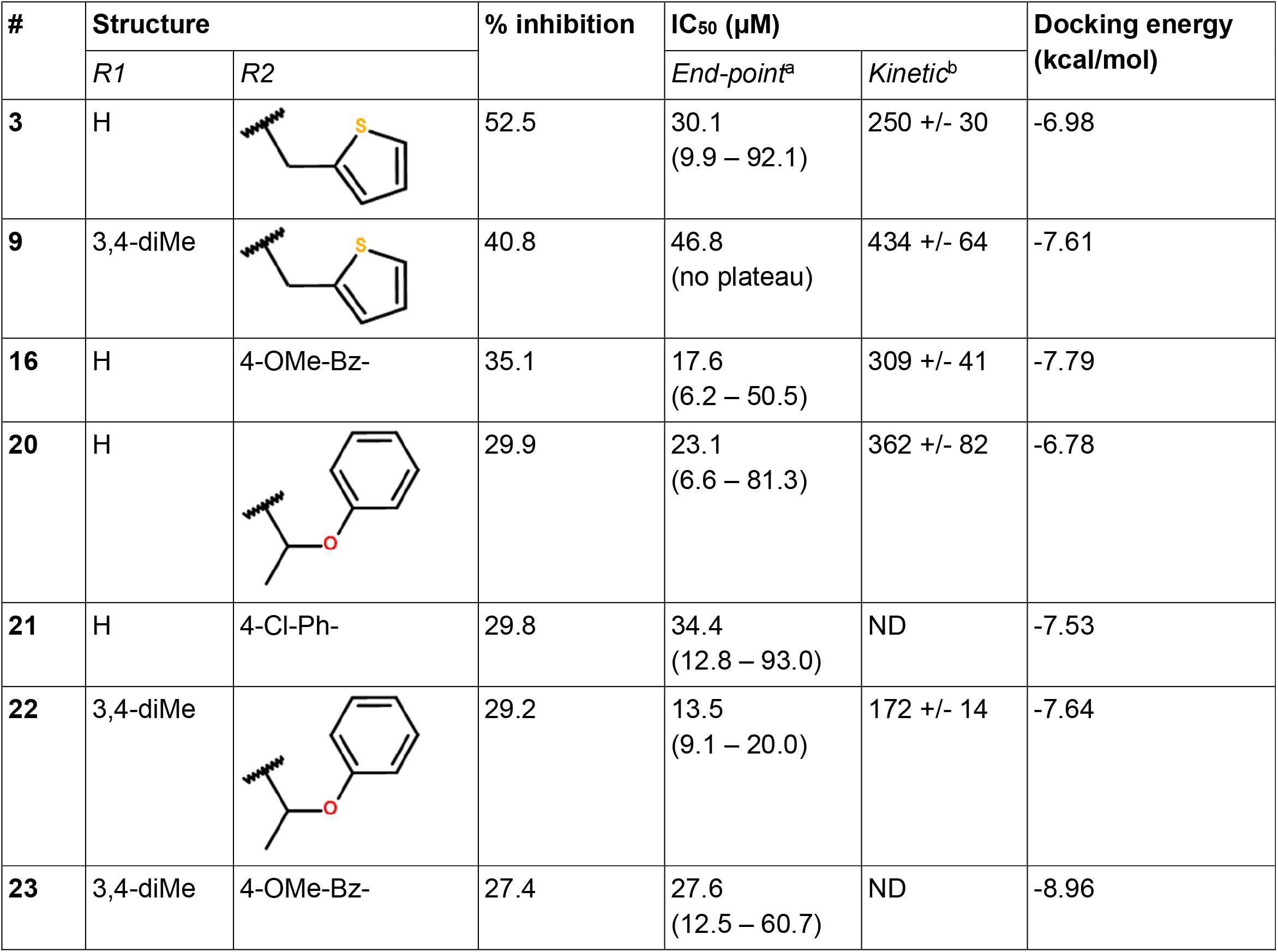
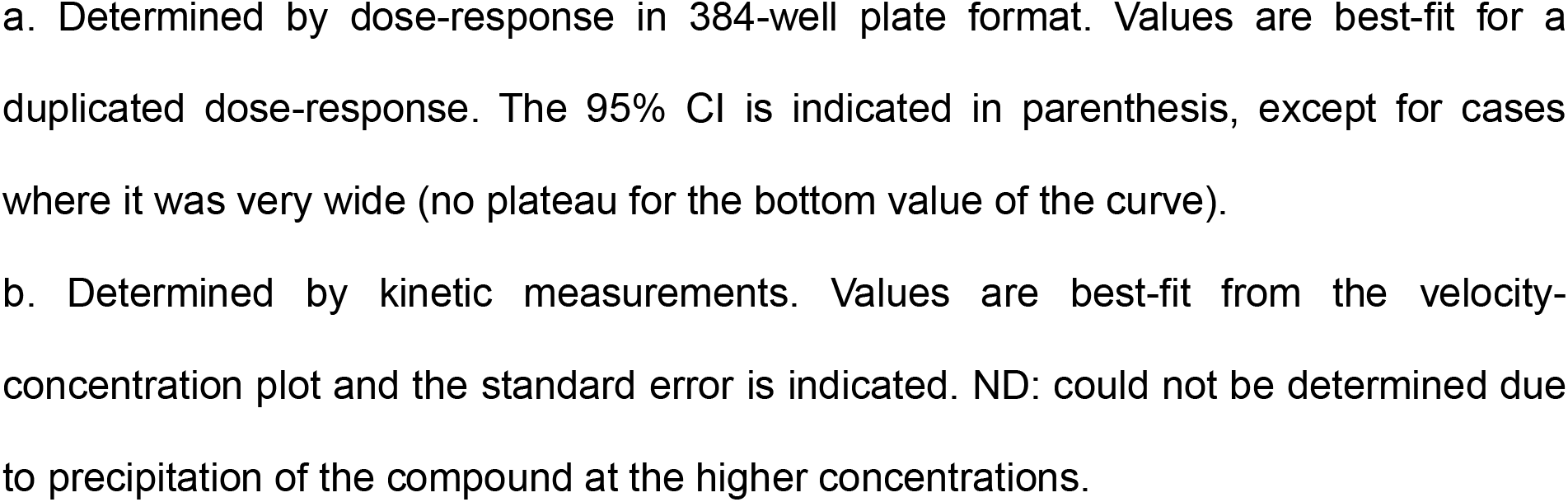
Activity profile of the compounds from cluster II.

| # | Structure |  | % inhibition | IC<sub>50</sub> (μM) |  | Docking energy (kcal/mol) |
| --- | --- | --- | --- | --- | --- | --- |
|  | R1 | R2 |  | End-point<sup>a</sup> | Kinetic<sup>b</sup> |  |
| 3 | H |  | 52.5 | 30.1<br>(9.9 – 92.1) | 250 +/- 30 | -6.98 |
| 9 | 3,4-diMe |  | 40.8 | 46.8<br>(no plateau) | 434 +/- 64 | -7.61 |
| 16 | H | 4-OMe-Bz- | 35.1 | 17.6<br>(6.2 – 50.5) | 309 +/- 41 | -7.79 |
| 20 | H |  | 29.9 | 23.1<br>(6.6 – 81.3) | 362 +/- 82 | -6.78 |
| 21 | H | 4-Cl-Ph- | 29.8 | 34.4<br>(12.8 – 93.0) | ND | -7.53 |
| 22 | 3,4-diMe |  | 29.2 | 13.5<br>(9.1 – 20.0) | 172 +/- 14 | -7.64 |
| 23 | 3,4-diMe | 4-OMe-Bz- | 27.4 | 27.6<br>(12.5 – 60.7) | ND | -8.96 |

a. Determined by dose-response in 384-well plate format. Values are best-fit for a duplicated dose-response. The 95% CI is indicated in parenthesis, except for cases where it was very wide (no plateau for the bottom value of the curve). b. Determined by kinetic measurements. Values are best-fit from the velocity-concentration plot and the standard error is indicated. ND: could not be determined due to precipitation of the compound at the higher concentrations.

Because *Mtb*FolB is able to transform the initial substrate into 3 different species (including 2 reversibly) [21], complex experiments would be needed to conclude on the inhibition mechanism. Here, we assumed that the compounds could bind within the active site (formed at the junction between 2 monomers of *Mtb*FolB) and conducted docking studies to explore their mode of binding. The estimated free energies of the top docking poses are indicated in **Table 2**. Interestingly, not all 7 derivatives adopted a similar pose and two distinct modes of binding were observed. In the first pose (compounds **9**, **16**, **22** and **23**), the carbonyl of the pyrazol-3-one core is engaged in hydrogen-bonding with the tyrosine and lysine of the active site (Tyr54D, Lys99A; A, D refer to the *Mtb*FolB subunit: two are necessary to form an active site), as well as π-π stacking interactions with Tyr54 (**Figure 3A** and **Figure S5**). The N-carbonyl side-chains were found to interact either through hydrophobic contacts (**9**, **22**) or hydrogen-bonding (**16**, **23**) with nearby residues. In the second pose however (compounds **3**, **20** and **21**), only the N-carbonyl side-chains were found to interact with the Tyr54D of the active site, through π-π stacking interactions (**Figure 3B** and **Figure S5**). Although the first pose is likely to induce inhibition of the catalytic activity, the second pose appears to have less affinity for the catalytic residues, which may lead to an easy displacement of the compound by the natural substrate. The existence of this second pose, less productive, may explain the low inhibition potential of the compounds from this cluster. Indeed, no compound in this series showed inhibitory activity against *M. tuberculosis* (data not shown).

**Figure 3.**
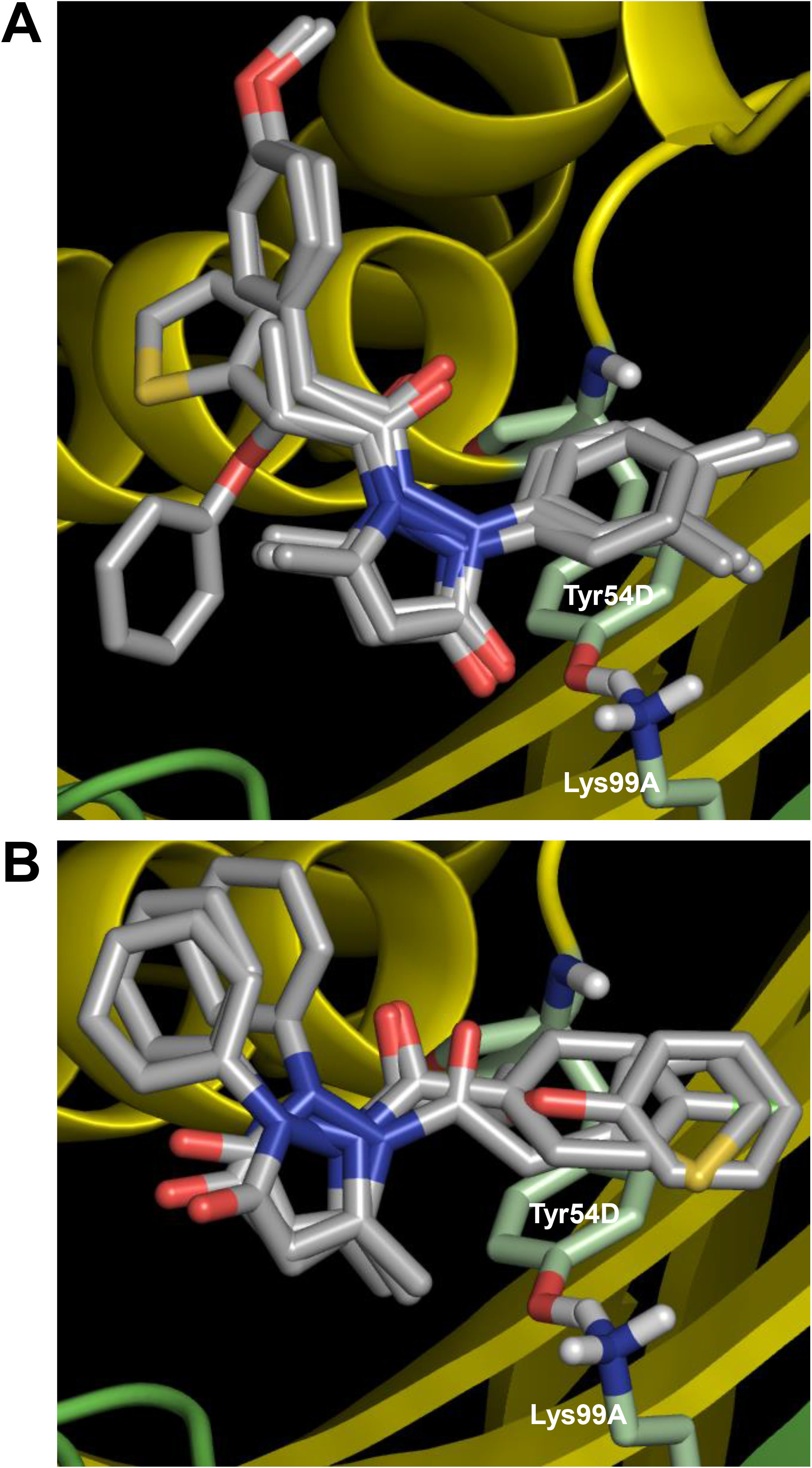
Docking results for compounds of cluster II. **A**. First docking pose, obtained with compounds **9**, **16**, **22** and **23**. The carbonyl of the pyrazol-3-one core is engaged in hydrogen-bonding with the tyrosine and lysine of the active site (Tyr54D, Lys99A; A, D refer to the *Mtb*FolB subunit: two are necessary to form an active site), as well as π-π stacking interactions with Tyr54D. **B**. Second docking pose (compounds **3**, **20** and **21**), where only the N-carbonyl side-chains interact with the Tyr54D of the active site through π-π stacking interactions. For both panels, compound backbones are indicated in gray, with nitrogen in blue, sulfur in light orange and oxygen in red. Residues of the active site are indicated in light green and labelled in white, with nitrogen in blue, hydrogen in light gray and oxygen in red. The two *Mtb*FolB monomers required to compose the active site are shown as yellow or green ribbon. PDB: 1NBU.

### Sulfonamide series (Cluster III)

The second series included 2 promising derivatives, namely **5** and **13**, displaying IC_50_ values of 2.6 and 15.4 μM, respectively. Considering the relative ease of access to these compounds and other derivatives (1 step reaction with the appropriate reagents), we decided to draw a preliminary structure activity profile of this series of compounds. In a first step, we explored the effect of varying the substituent connected to the sulfur atom of the sulfonamide (S-series, **Table 3**). Our results indicated that the activity could be retained for a variety of substituents. In particular, the pyrazole **25** and the isoxazole **27** both showed good activities (IC_50_ values of 6.8 μM and 9.2 μM, respectively) and the longer, flexible diphenyl ether chain of **30** could also be tolerated, although the resulting inhibition was more modest (IC_50_ value of 31.7 μM). In a second step, we explored the substituents on the amide side (N-series) and noted that variations at this position might be more constrained (**Table 4**). While the piperidine **32** had slightly better activity compared to the reference secondary amine (IC_50_ value of 11.3 μM against 15.4 μM), the primary amine **31** was less active (IC_50_ value of 37.8 μM) and the imidazole **33** together with the amide **34** were both inactive. Substituting the piperidine led to either decrease (**35**, **36**) or loss (**38**) of the activity. Replacing the secondary amine with an isopropyl also led to the loss of activity, as seen in compound **37**. Interestingly however, the phenyl piperazine-linked bi-sulfonamide **39** maintained a good activity (IC_50_ value of 10.6 μM), indicating that this type of bi-sulfonamide could present a viable strategy for further optimizations.

**Table 3.**
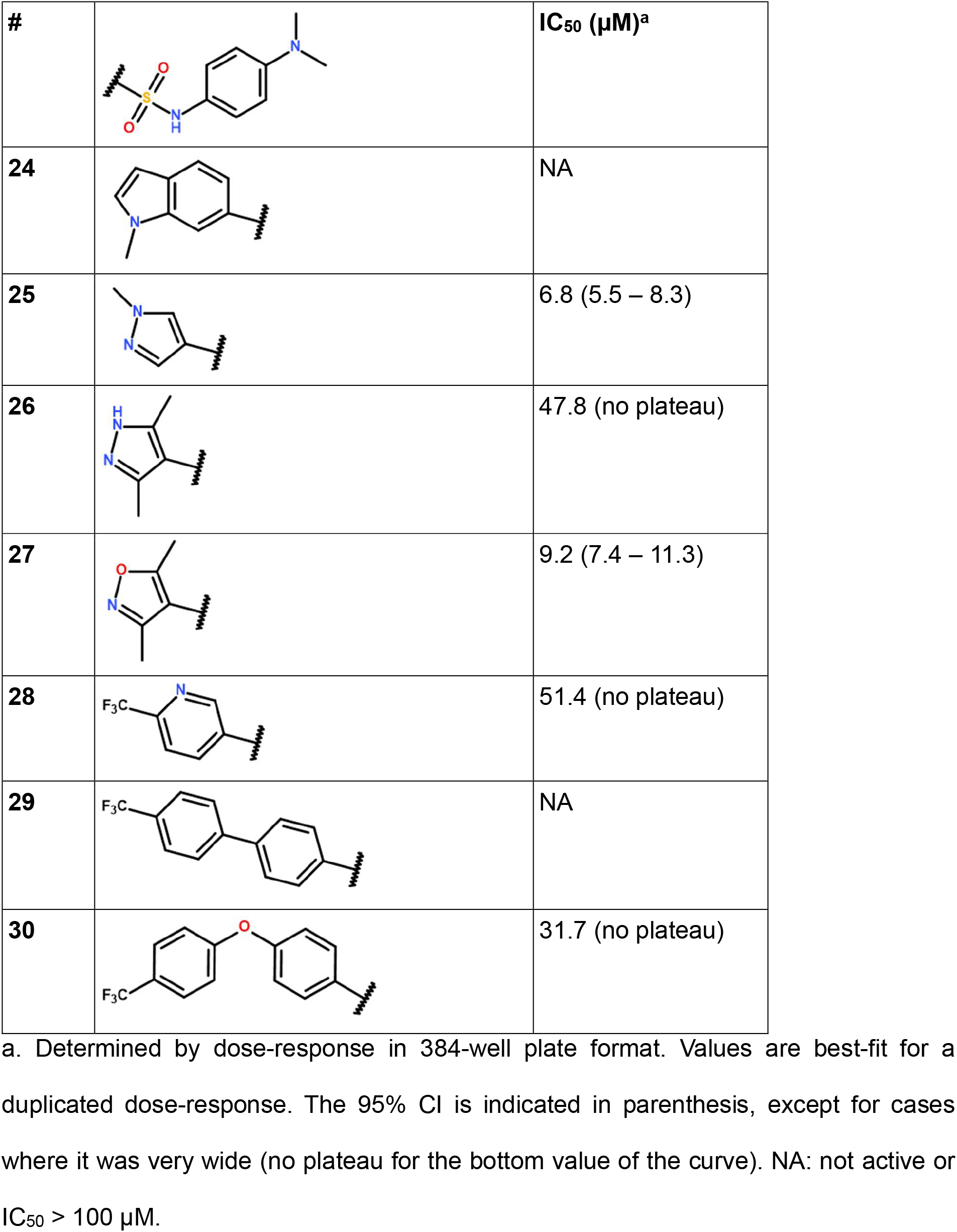
Activity profile of the derivatives from cluster III (S-series)

| # |  | IC<sub>50</sub> (μM)<sup>a</sup> |
| --- | --- | --- |
| 24 |  | NA |
| 25 |  | 6.8 (5.5 – 8.3) |
| 26 |  | 47.8 (no plateau) |
| 27 |  | 9.2 (7.4 – 11.3) |
| 28 |  | 51.4 (no plateau) |
| 29 |  | NA |
| 30 |  | 31.7 (no plateau) |

a. Determined by dose-response in 384-well plate format. Values are best-fit for a duplicated dose-response. The 95% CI is indicated in parenthesis, except for cases where it was very wide (no plateau for the bottom value of the curve). NA: not active or IC<sub>50</sub> > 100 μM.

**Table 4.**
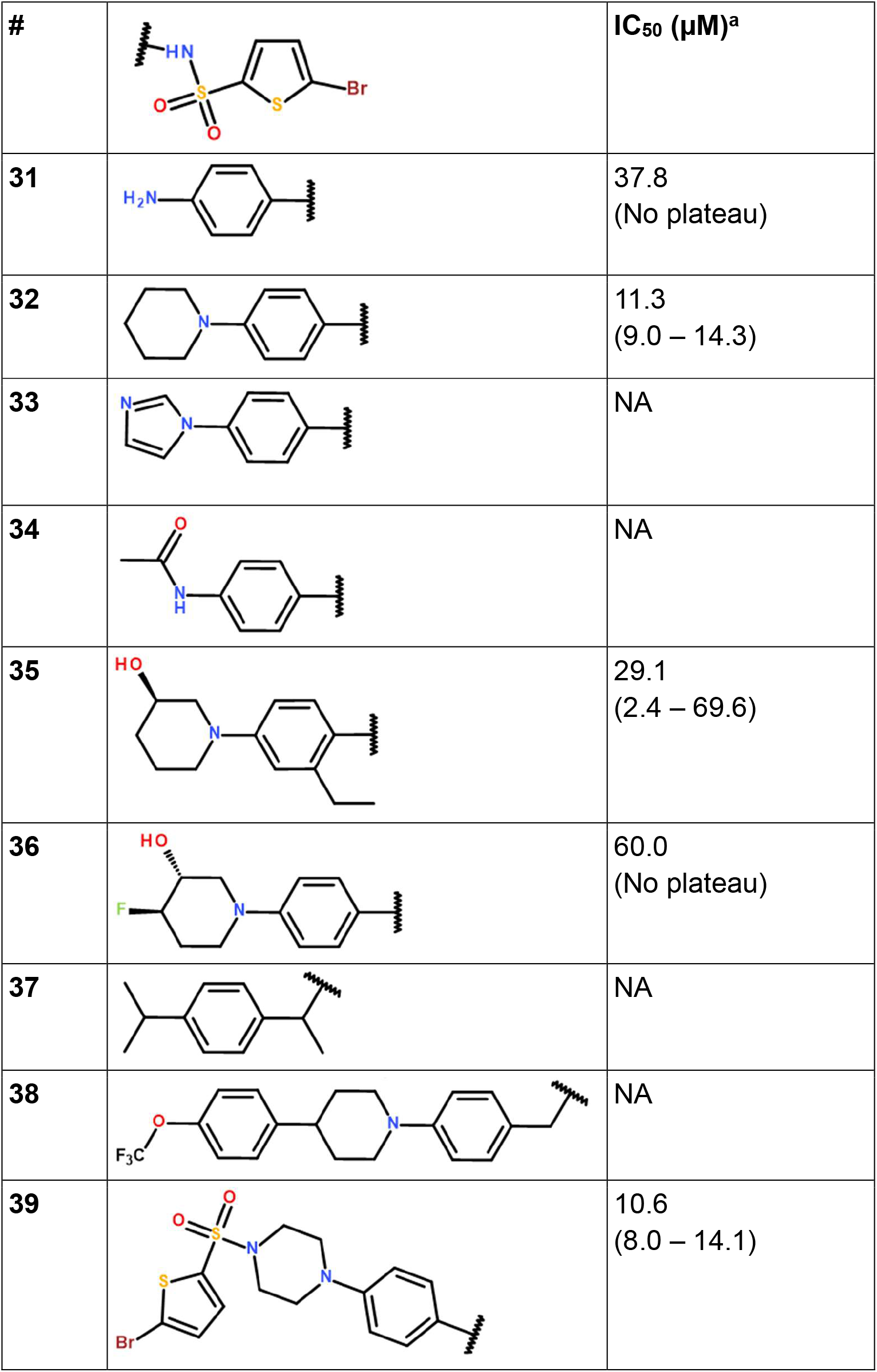
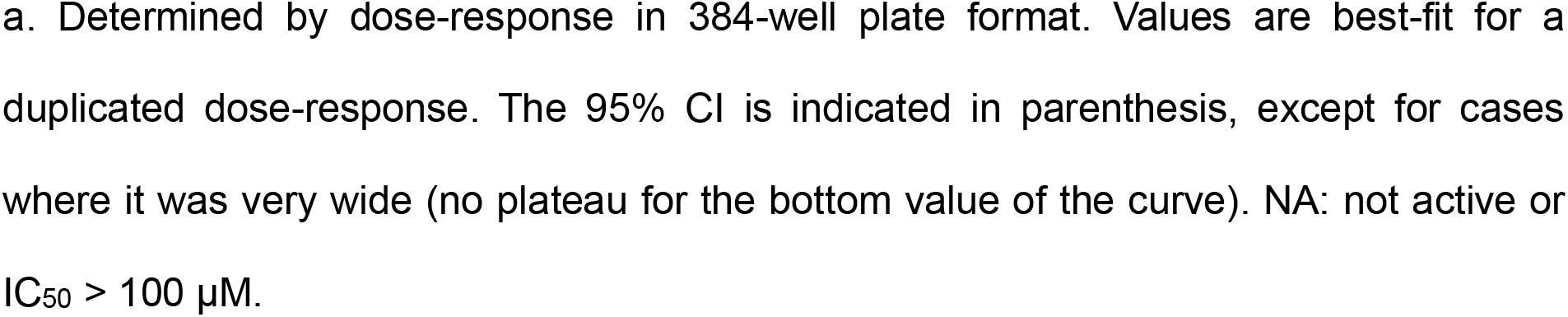
Activity profile of the derivatives from cluster III (N-series)

| # |  | IC<sub>50</sub> (μM)<sup>a</sup> |
| --- | --- | --- |
| 31 |  | 37.8<br>(No plateau) |
| 32 |  | 11.3<br>(9.0 – 14.3) |
| 33 |  | NA |
| 34 |  | NA |
| 35 |  | 29.1<br>(2.4 – 69.6) |
| 36 |  | 60.0<br>(No plateau) |
| 37 |  | NA |
| 38 |  | NA |
| 39 |  | 10.6<br>(8.0 – 14.1) |

a. Determined by dose-response in 384-well plate format. Values are best-fit for a duplicated dose-response. The 95% CI is indicated in parenthesis, except for cases where it was very wide (no plateau for the bottom value of the curve). NA: not active or $IC_{50} > 100 \mu M$ .

Similarly to the compounds of the pyrazol-3-one series (Cluster II), kinetic and time-dependent inhibition studies were initiated for the sulfonamide series. Despite reasonable computed logP and polar surface area (TPSA) values (< 3 and < 120, respectively; **Table S1**), solubility issues appeared for most compounds of this family preventing studies at concentrations exceeding 20 to 40 µM, which was insufficient to complete the required experiments. We thus decided to focus instead on docking studies in order to gain more insight into the potential mode of binding of these compounds in the active site of *Mtb*FolB. We chose to include compounds **5**, **13**, **27**, **32** and **39** as they were among the most active derivatives identified for this series. Compounds **24** and **34**, which were found to be inactive, were also included for comparison. As shown in **Figure 4A**, compounds **13**, **32** and **39** aligned on the same pose within the active site and displayed several interactions with the tyrosine and lysine of the active site (Tyr54D, Lys99A), through π-π stacking interactions with Tyr54D and both the benzyl and thiophene rings on each side of the sulfonamide, as well as hydrogen-bonding between the Lys99A and the oxygen of the sulfonamide (distance ∼3.0 Å). While the S-moiety of the sulfonamide lied within the catalytic pocket, the N-moiety rested at the entrance of the cavity, lined by alkyl residues Val55D and Ile104A (**Figure 4B**). Compound **27** had a similar orientation but displayed a different pose in the active site, so that the Lys99A was apparently not engaged in the interaction (**Figures S6A, S7**). Compound **5** in contrast showed an opposite orientation, with the S-moiety pointing towards the outside, similarly to the two inactive compounds **24** and **34** (**Figure S6**). The presence of two hydrophobic and bulky amino-acids at the entrance of the active site (Val55D and Ile104A) may explain the lower tolerance for variation on this side of the molecule, in particular the loss of activity seen for the quite rigid and polar imidazole **33** and the rigid and bulky 4-isopropylbenzyl **37**. It is more complicated however to understand why the amine **31** and amide **34** are inactive solely based on these docking studies. Molecular dynamic experiments would be required to better appreciate the interactions within the active site. Other parameters might also have to be considered to explain this lack of activity, like non-specific binding elsewhere in the *Mtb*FolB tetramer prior to the formation of the octamer, which is expected to occur upon substrate binding [24].

**Figure 4.**
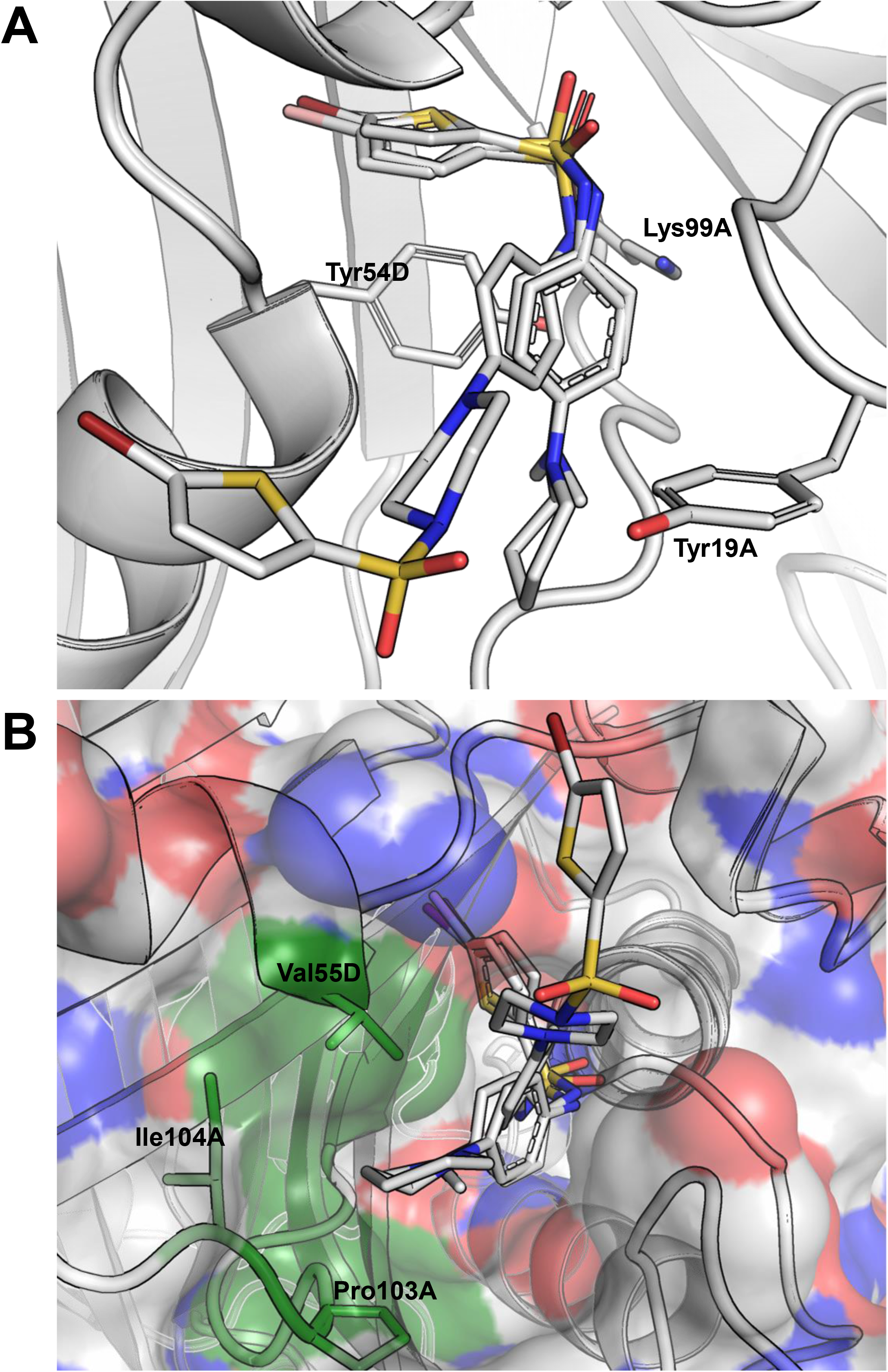
Docking results for compounds **13**, **32** and **39** of cluster III. **A**. Docking pose obtained for the three compounds, where they are seen interacting with the tyrosine of the active site (Tyr54D; A, D refer to the *Mtb*FolB subunit: two are necessary to form an active site) through π-π stacking interactions with both the benzyl and thiophene rings on each side of the sulfonamide. Hydrogen-bonding is also possible between the lysine of the active site (Lys99A) and the oxygen of the sulfonamides (distance ∼3.0 Å). Compound backbones are indicated in gray, with nitrogen in blue, sulfur in light orange and oxygen in red. PDB: 1NBU. **B**. Surface view of the *Mtb*FolB dimer, showing that in this docking pose the S-moiety of these sulfonamides lye within the catalytic pocket, while the N-moieties rest at the entrance of the cavity, lined by residues Val55D, Ile104A and Pro103A. Surface transparency is 20% and colors are gray for carbon, blue for nitrogen and red for oxygen. The three residues lining the cavity entrance are labelled and colored in green. PDB: 1NBU.

Interestingly, compound **13** showed a modest inhibitory activity against the bacillus, with a MIC of 55.4 µM. To our knowledge, this is the first compound directed against *Mtb*FolB that is also found to display an antimycobacterial activity. This finding paves the way for further medicinal chemistry efforts to improve the antimycobacterial potency of this lead compound **13**.

## Acknowledgments

Institut Pasteur Korea is a member of the Institut Pasteur International Network (https://pasteur-network.org/). This work was supported by the national research foundation of Korea (NRF), funded by the Korean ministry of science, ICT and technology (MSIT, project numbers NRF-2017M3A9G6068246 and NRF-2019R1A2C2084652) as well as the Gyeonggi province, Korea. VD was supported by the French ministry of foreign affairs. This work was also supported by the National Institute of Science and Technology on Tuberculosis (CNPq-FAPERGS-CAPES) [grant number 421703–2017-2 and 17– 1265-8]. C. V. Bizarro, L. A. Basso, and P. Machado are Research Career Awardees of the National Research Council of Brazil (CNPq). This study was financed in part by the Coordenação de Aperfeiçoamento de Pessoal de Nível Superior, Brazil (CAPES), Finance Code 001.

## Notes

### Competing Interest Statement

The authors have declared no competing interest.

