## Supplemental Figures for "Identification and characterization of new structural scaffolds modulating the activity of *Mycobacterium tuberculosis* dihydroneopterin aldolase (FolB) *in vitro*"

#### Current address: R&D Center, Insol Co., Ltd., Hanam, Gyeonggi, Republic of Korea

**Keywords**

### Supplementary figures

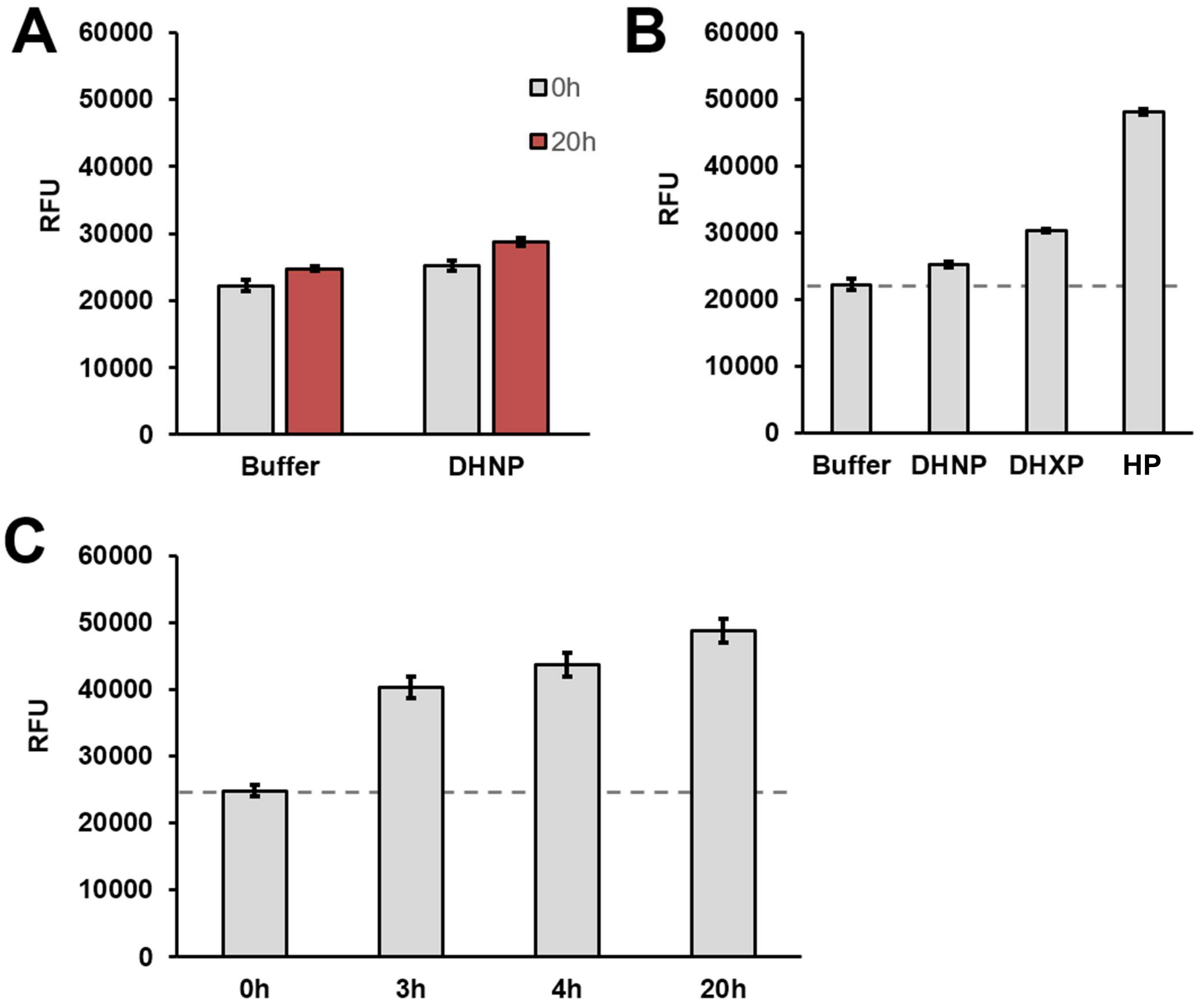


**Figure S1. Optimization of a fluorometric assay for the estimation of FolB activity**. **A**. Evolution of the fluorescence of the buffer solution used for the reaction (25 mM Tris-HCl pH 8.0, 50 mM NaCl, 5% glycerol), alone (buffer), and supplemented with 5 µM of FolB substrate (DHNP), after an incubation period of 20h at room temperature. **B**. Fluorescence of the substrate (DHNP), thermodynamic (DHXP), and kinetic (HP) products of the FolB reaction when solubilized in the buffer at a concentration of 5 µM. **C**. End-points fluorescence measurement of the solution obtained after incubation of FolB and its substrate (DHNP, 5 µM) for the indicated period at room temperature. For all experiments, fluorescence was recorded using Ex. 405 nm and Em. 535 nm. A dashed gray line is used to indicate the average fluorescence of the substrate alone (background). Data shown are average and SD for at least 3 technical replicates.

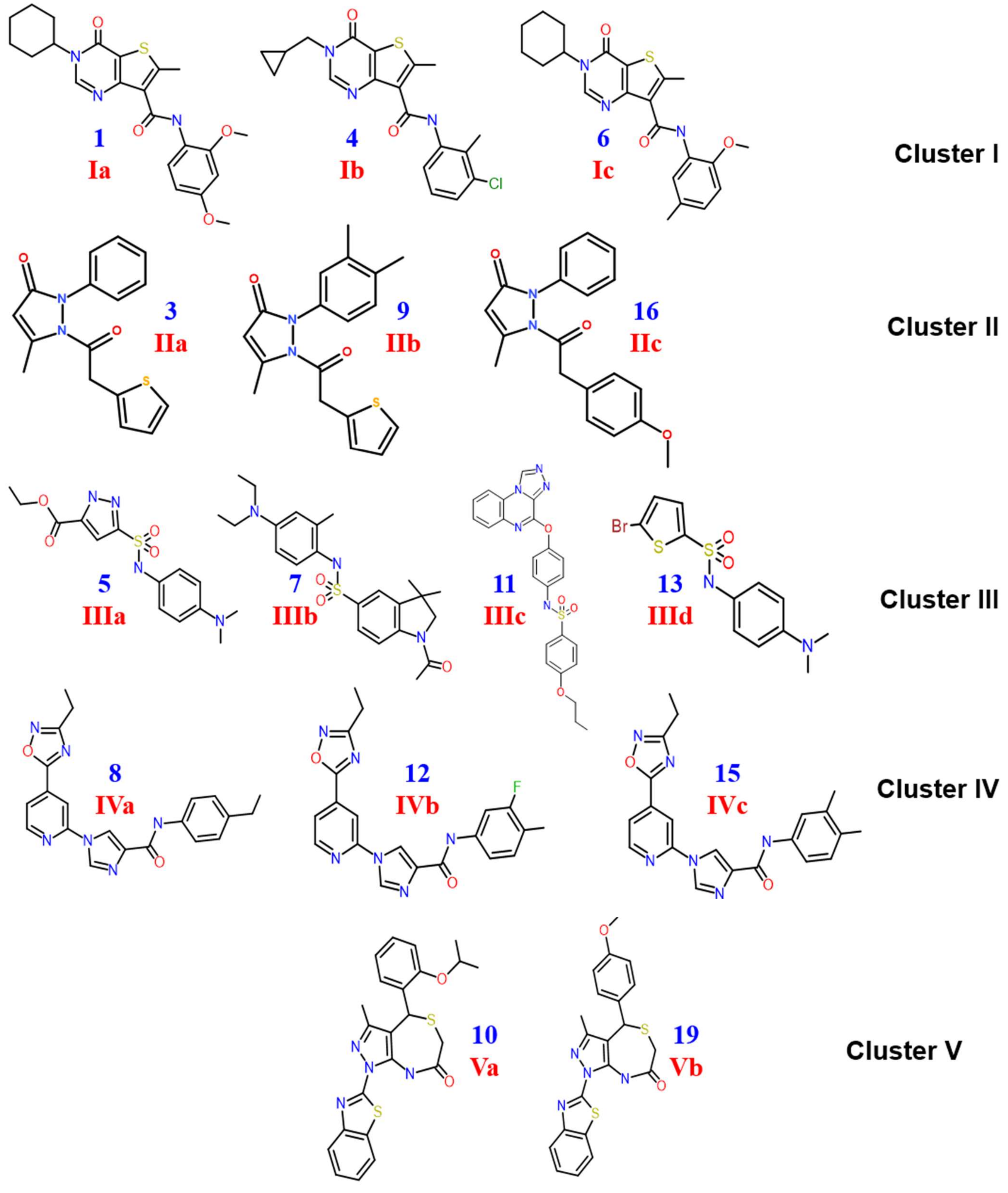
1
Ia
4
Ib
6
Ic
Cluster I
3
IIa
9
IIb
16
IIc
Cluster II
5
IIIa
7
IIIb
11
IIIc
13
IIId
Cluster III
8
IVa
12
IVb
15
IVc
Cluster IV
10
Va
19
Vb
Cluster V

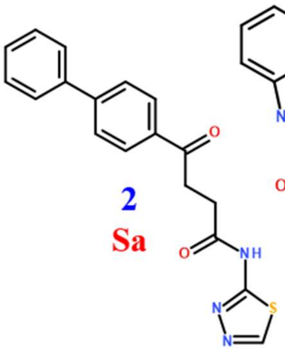

**2**
**Sa**

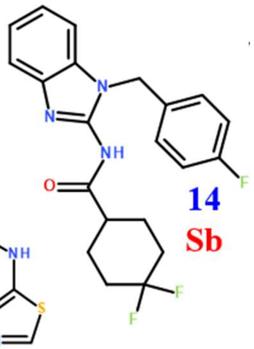

**14**
**Sb**

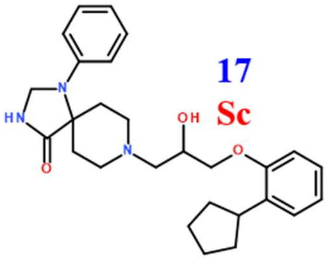

**17**
**Sc**

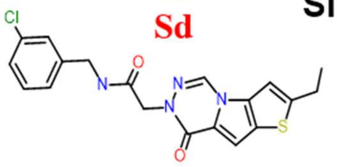

**18**
**Sd**

**Singletons**

**Figure S2. Structure of the 19 hits selected at a 30% inhibition threshold, arranged by cluster**. Blue numbering is as in **Table 1** from the main manuscript. Red numbering is as in the chemistry section of the supplementary methods.

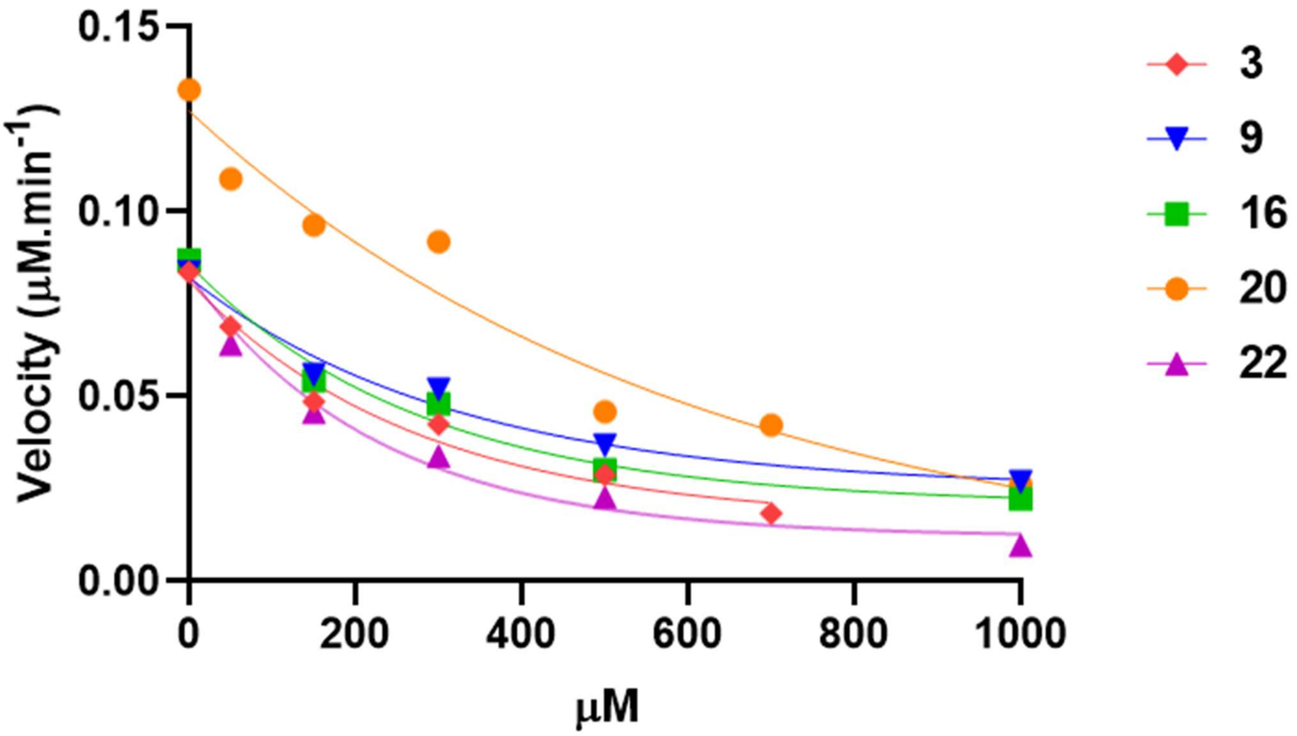


**Figure S3. Kinetic inhibition studies for the pyrazol-3-one series (Cluster II)**. Each reaction velocity (μM.min$^{-1}$) was calculated by linear regression of a 6-min kinetic (20 data points) as described in the Methods of the main manuscript. Data shown are for a single, representative experiment. Compounds **3**, **9** and **16** are the screening hits confirmed by dose-response. Compounds 20 and 22 are the two additional derivatives synthesized with the highest inhibition activity, as per the dose-response assay (see **Table 2**).

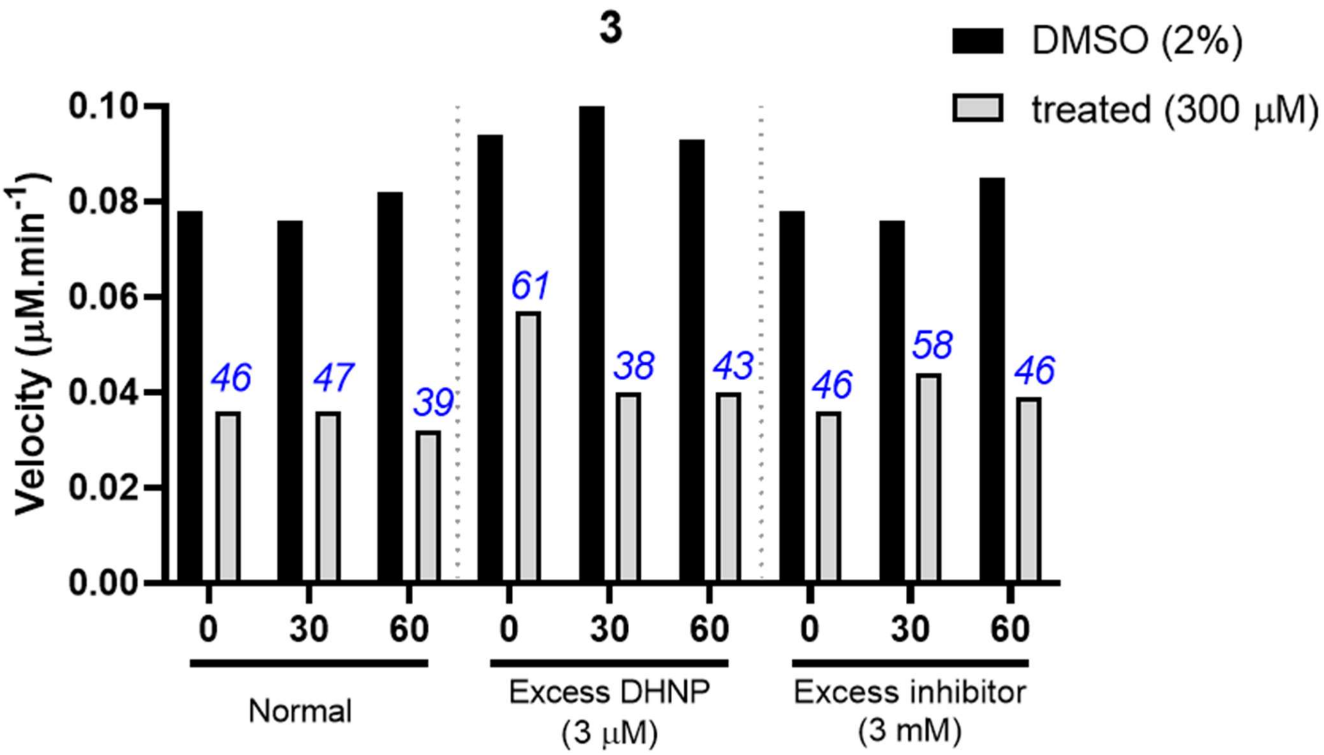
3
DMSO (2%)
treated (300 μM)
Velocity (μM.min-1)
0.10
0.08
0.06
0.04
0.02
0.00
46
47
39
61
38
43
46
58
46
0
30
60
0
30
60
0
30
60
Normal
Excess DHNP
(3 μM)
Excess inhibitor
(3 mM)

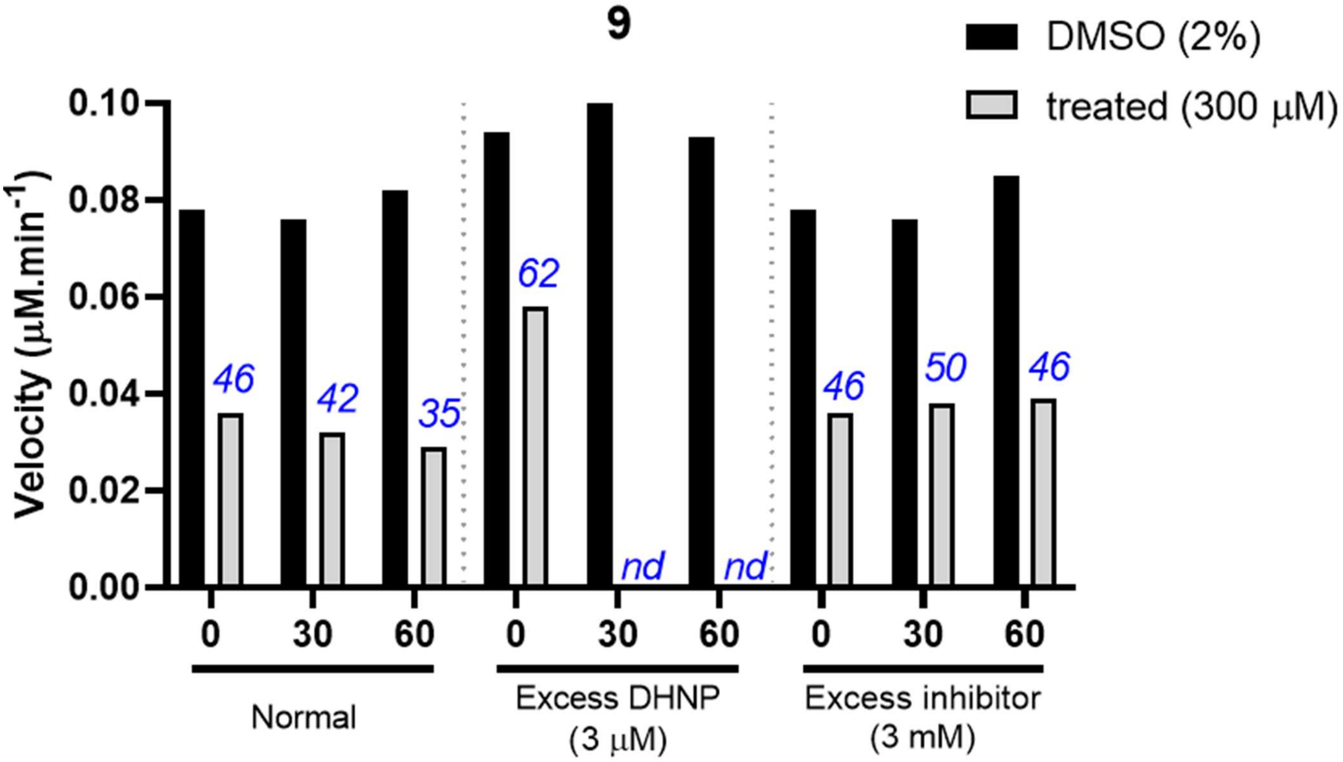
9
DMSO (2%)
treated (300 μM)
Velocity (μM.min-1)
0.10
0.08
0.06
0.04
0.02
0.00
46
42
35
62
nd
nd
46
50
46
0
30
60
0
30
60
0
30
60
Normal
Excess DHNP
(3 μM)
Excess inhibitor
(3 mM)

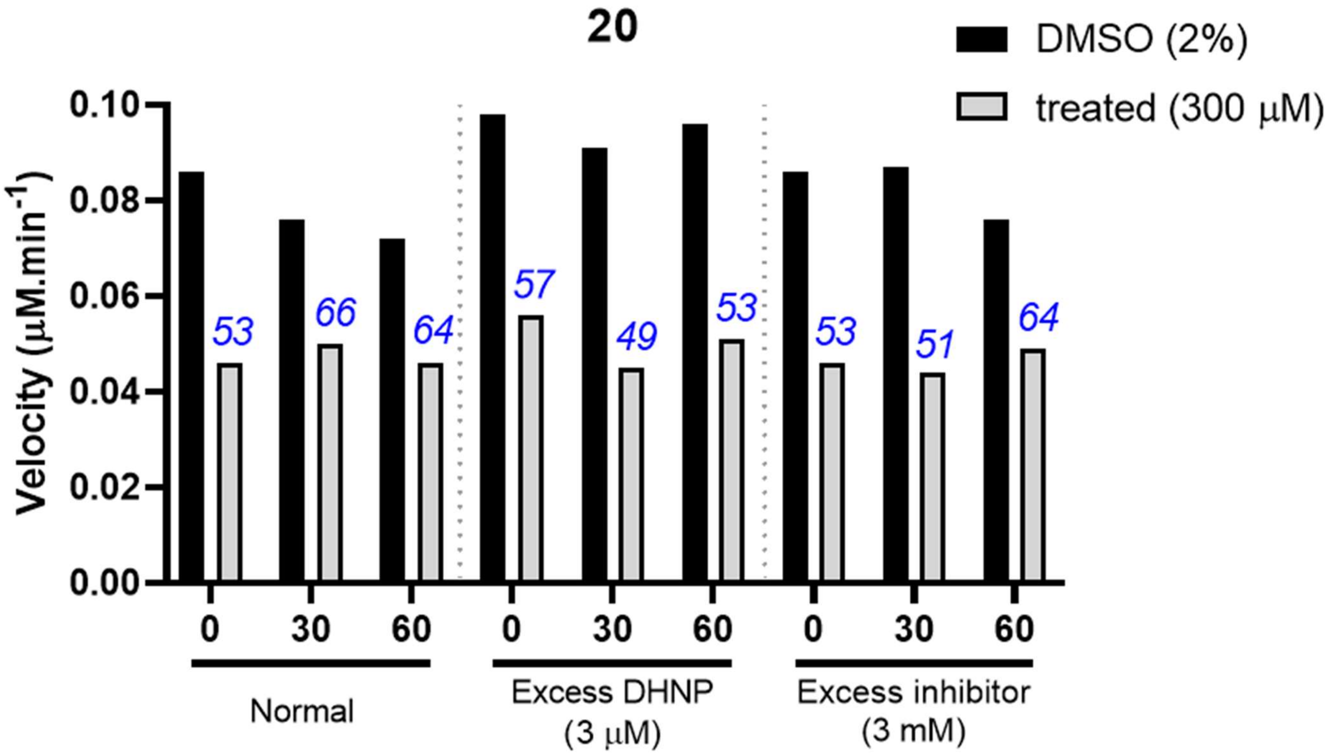


**Figure S4. Time-dependent inhibition by pyrazol-3-one series (Cluster II)**. *Mtb*FolB (300 nM) was incubated at room temperature in the reaction buffer together with the compound (3, 9 or 20) prior to determination of the reaction velocity by fluorometry. At the corresponding time-point (0, 30 and 60 min), the substrate (DHNP) was added and the enzyme velocity determined by kinetic assay (see methods in main text). Different incubation and test conditions were used. *i*) Normal condition: *Mtb*FolB was incubated with the compound at 300 µM, then the velocity measured in presence of the compound (300 µM) and 1.5 µM of substrate. *ii*) Excess of substrate: *Mtb*FolB was incubated with the compound at 300 µM, then the velocity measured in presence of the compound (300 µM) and 3 µM (2×) of substrate. *iii*) Excess of inhibitor: *Mtb*FolB was incubated with the compound at 3 mM (10×), then the velocity measured in presence of the compound (300 µM) and 1.5 µM of substrate. For each sample, a control incubated and tested without any inhibitor was performed to obtain the $V_{MAX}$. Percentage of velocity obtained for the treated samples, as compared to the untreated ones, is indicated in blue of top of each column. *nd*: could not be determined due to compound precipitation. Each reaction velocity (µM.min$^{-1}$) was calculated by linear regression of a 6-min kinetic (20 data points). Data shown are for a single, representative experiment.

**9**

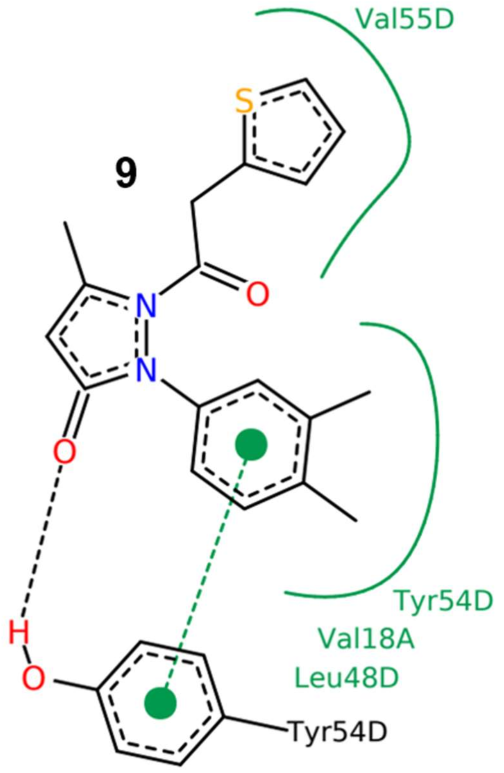
Val55D
S
N
N
O
O
Tyr54D
Val18A
Leu48D
H
O
Tyr54D

**22**

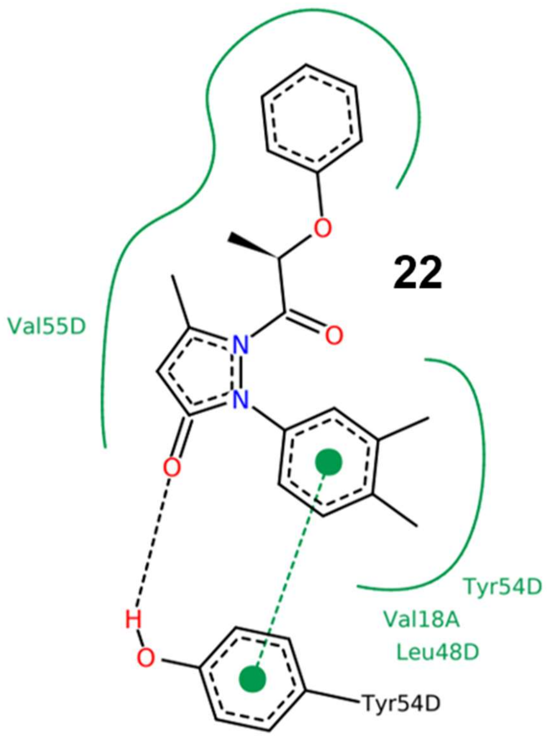
O
Val55D
N
O
N
O
Tyr54D
Val18A
Leu48D
H
O
Tyr54D

**16**

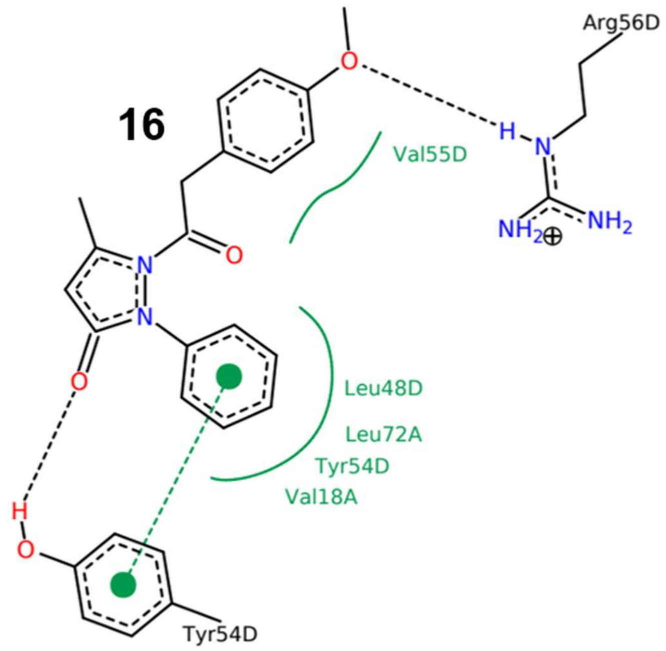
Arg56D
O
H
N
Val55D
NH2
NH2
N
O
N
O
Leu48D
Leu72A
Tyr54D
Val18A
H
O
Tyr54D

**23**

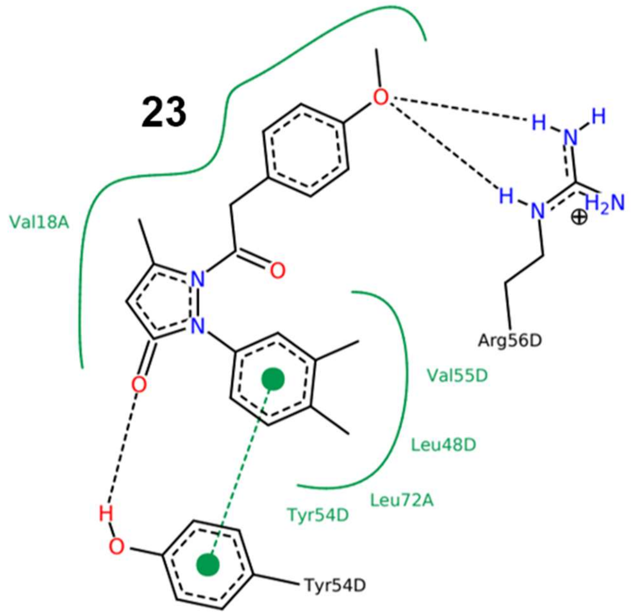
O
H
N
H
Val18A
H
N
H2N
N
O
N
Arg56D
Val55D
O
Leu48D
Leu72A
Tyr54D
H
O
Tyr54D

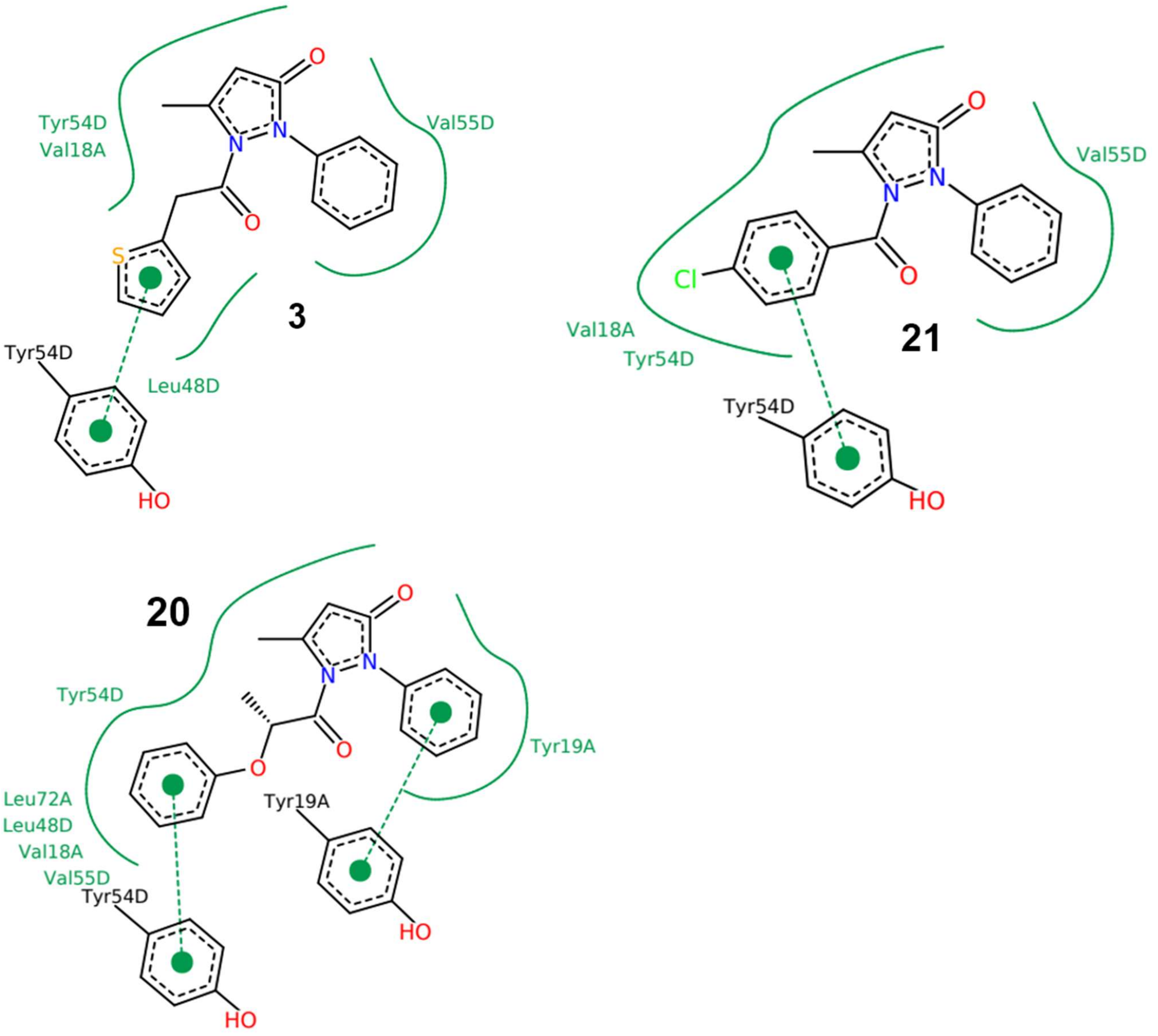


**Figure S5. 2D ligand interaction diagram obtained for the docking of compounds from the pyrazol-3-one series (Cluster II)**. Hydrophobic interactions are represented as green lines and π-π stacking interactions by green dashed lines. Images were generated with PoseView.

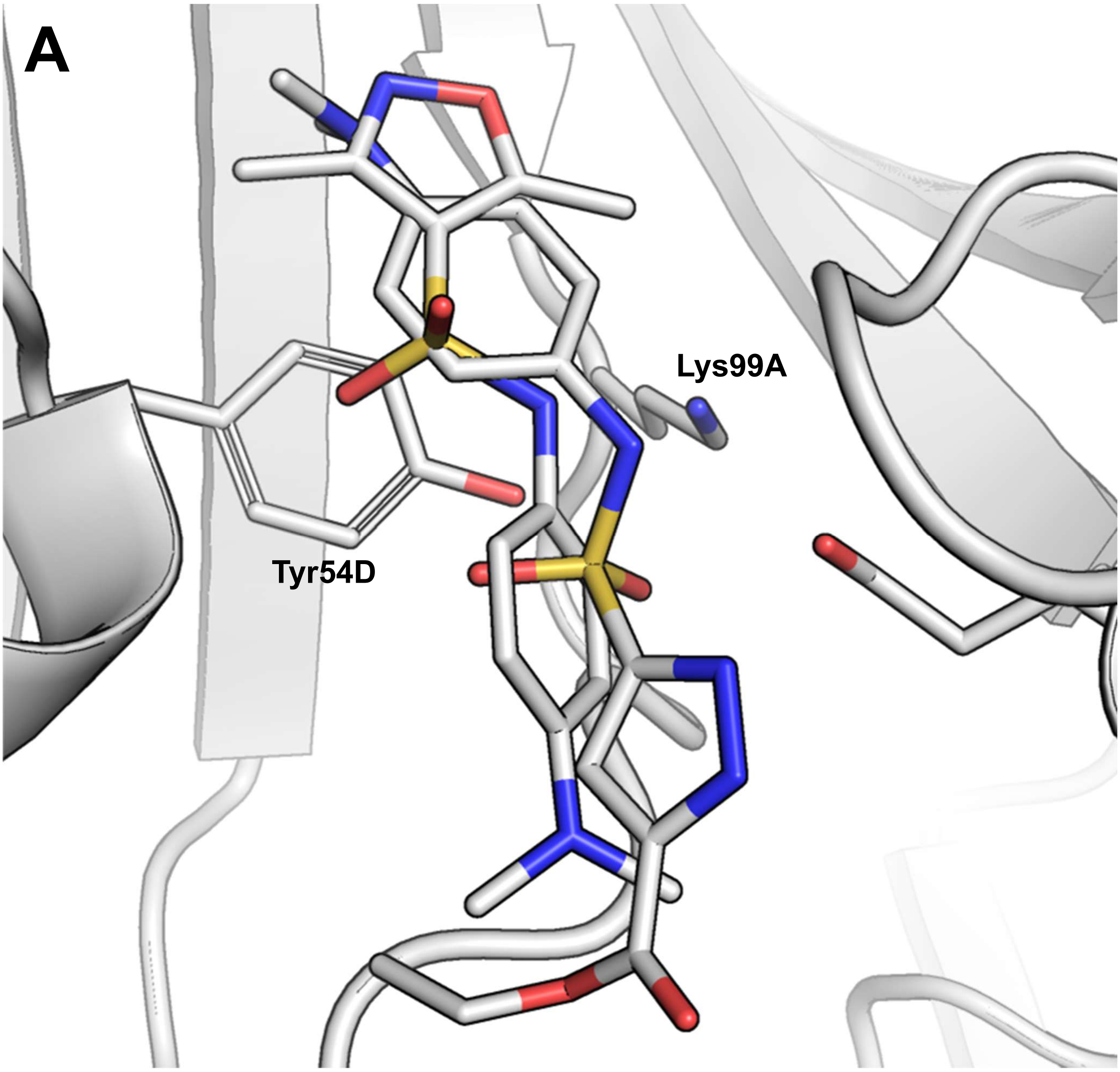
A
Lys99A
Tyr54D

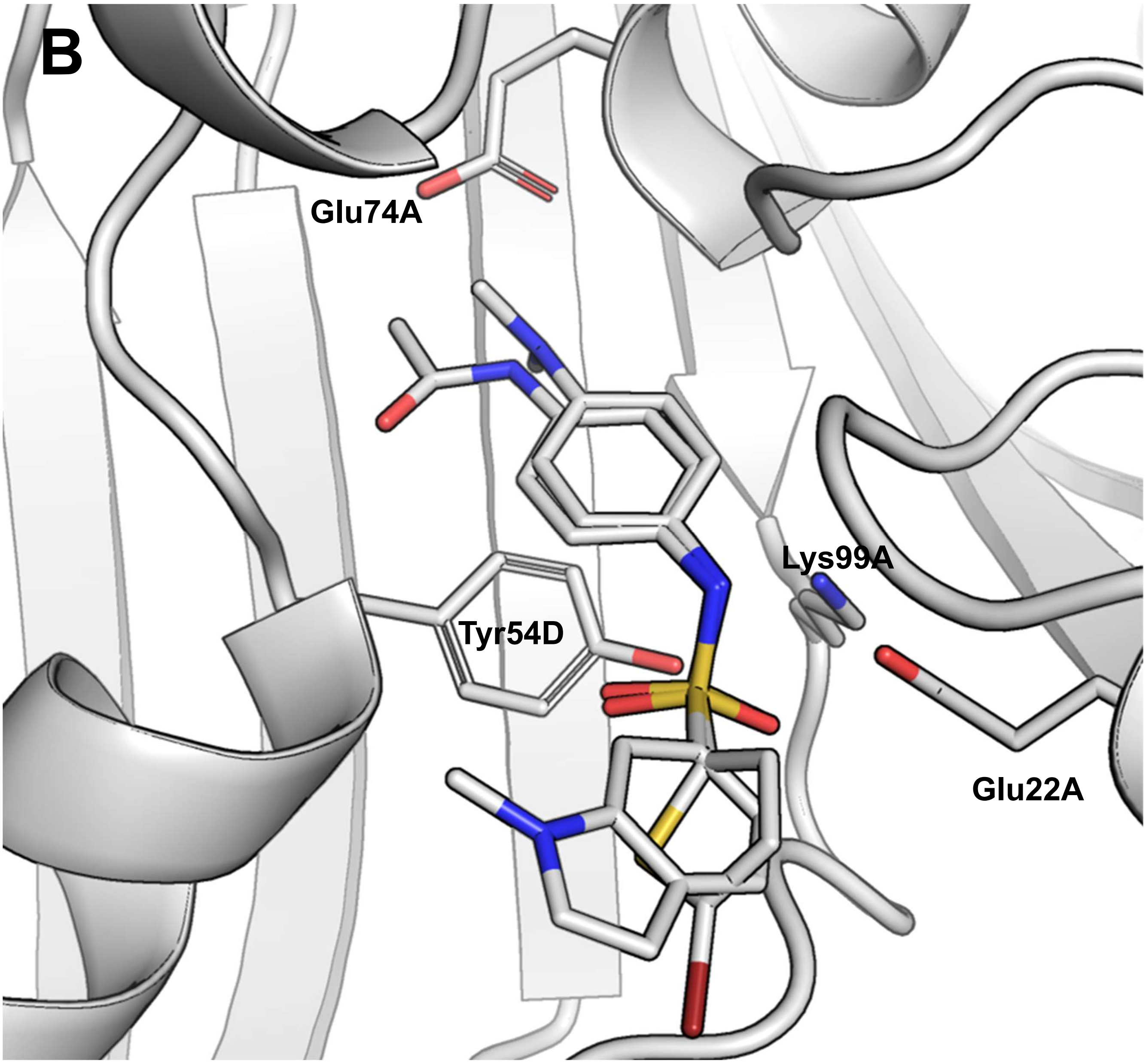


**Figure S6. Docking poses obtained for compounds of the sulfonamide series (Cluster III)**. **A**. Active compounds **5** and **27** are shown superimposed. **B**. Inactive compounds **24** and **34** are shown superimposed.

5

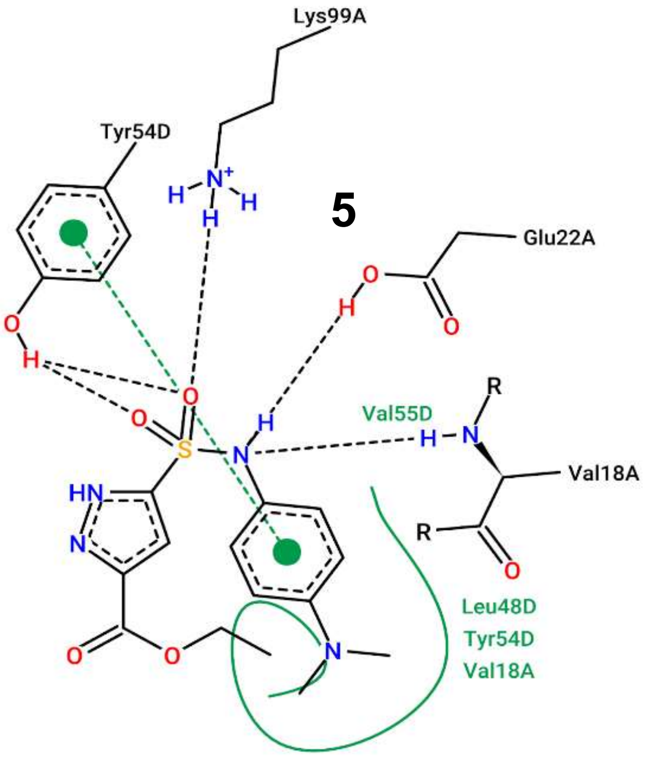
Lys99A
Tyr54D
Glu22A
Val55D
Val18A
Leu48D
Tyr54D
Val18A

13

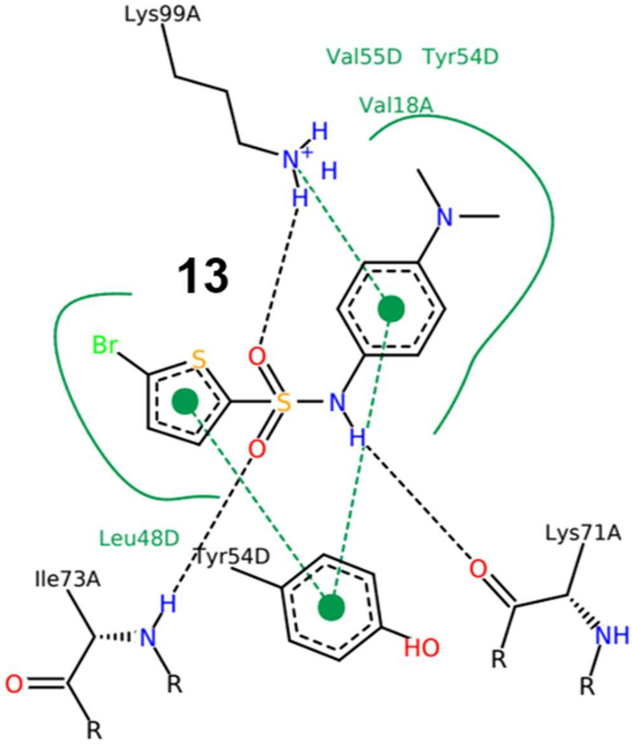
Lys99A
Val55D Tyr54D
Val18A
Br
Leu48D
Tyr54D
Ile73A
Lys71A

24

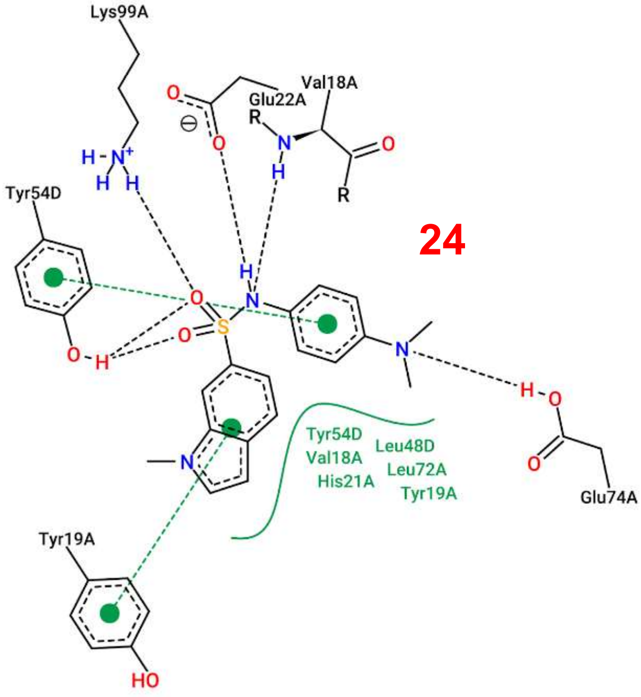
Lys99A
Val18A
Glu22A
Tyr54D
Tyr54D
Val18A
His21A
Leu48D
Leu72A
Tyr19A
Glu74A
Tyr19A

27

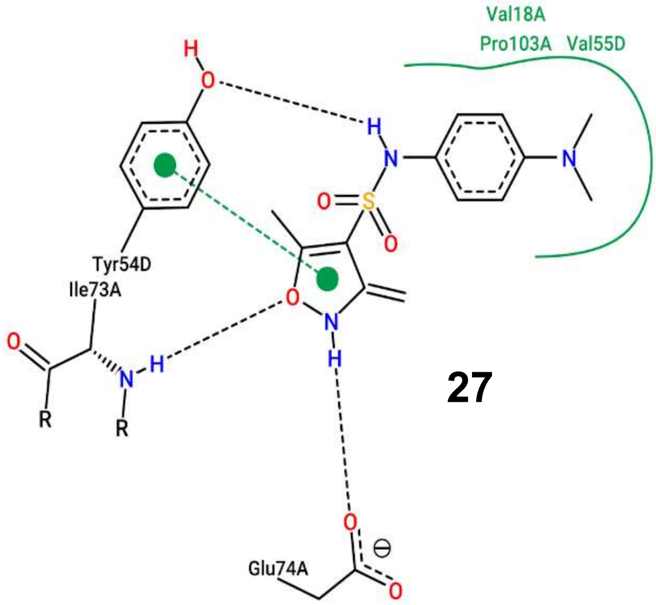
Val18A
Pro103A Val55D
Tyr54D
Ile73A
Glu74A

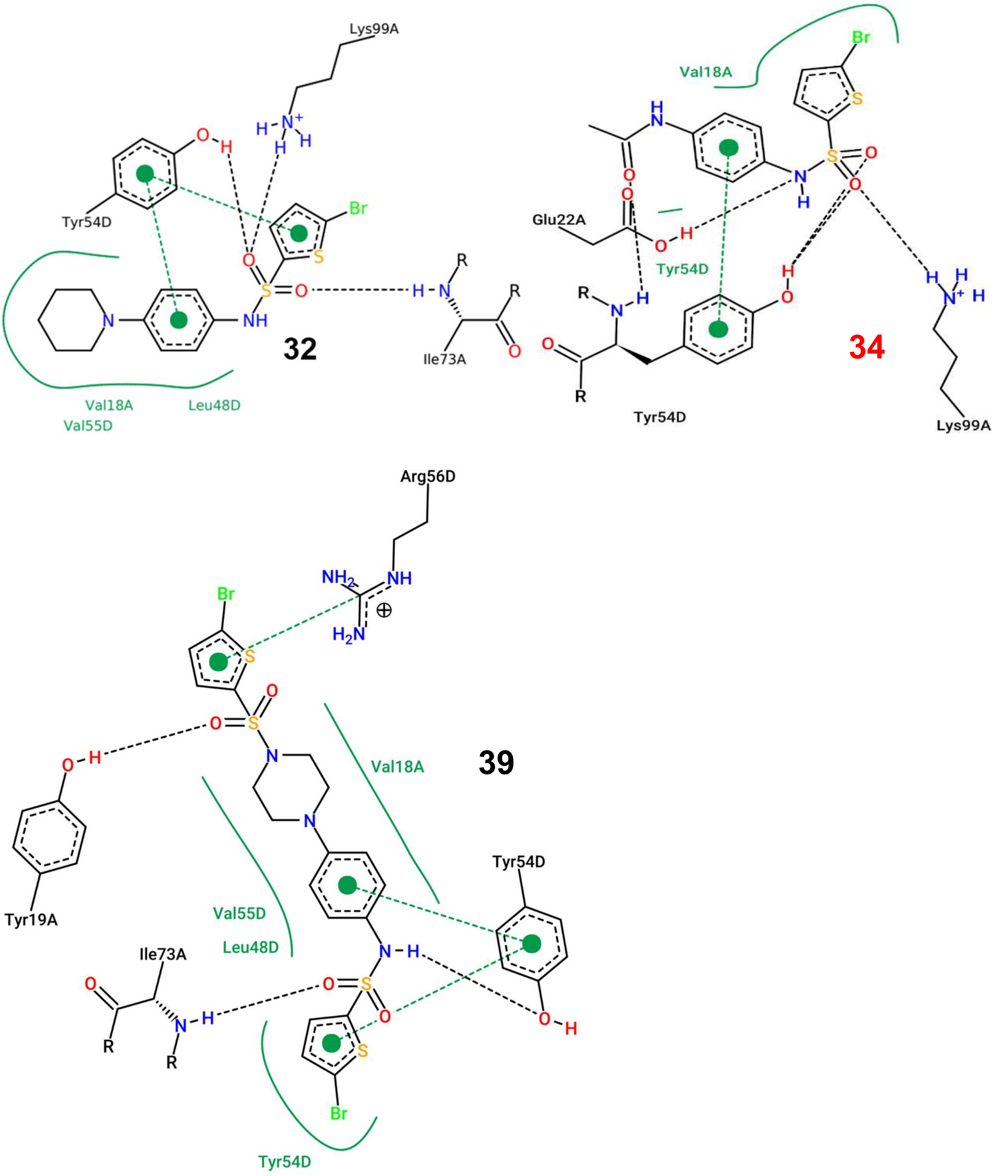


**Figure S7. 2D ligand interaction diagram obtained for the docking of compounds from the sulfonamide series (Cluster III)**. Compounds **24** and **34**, which were found to be inactive *in vitro* and were included for comparison purposes, are indicated in red. Hydrophobic interactions are represented as green lines, π-π stacking interactions by green dashed lines and hydrogen bonds by black dashed lines. Images were generated with PoseView.

**Table S1.** Computed hydrophobicity-related properties for the molecules of the sulfonamide cluster (Cluster III), obtained from the molinspiration web server (https://www.molinspiration.com). TPSA: topological polar surface area. Values < 3 for logP and < 120 for TPSA are usually considered acceptable for good solubility.

**A. Screening Hits**

| # | Structure | logP | TPSA |
|---|---|---|---|
| 5 | 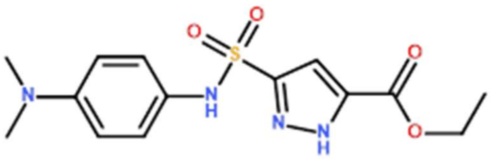 | 1.92 | 104.39 |
| 7 | 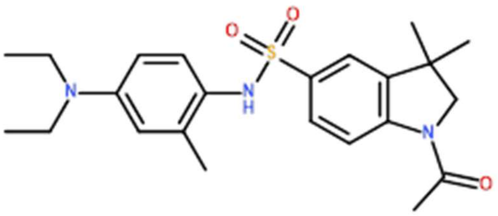 | 4.34 | 69.72 |
| 11 | 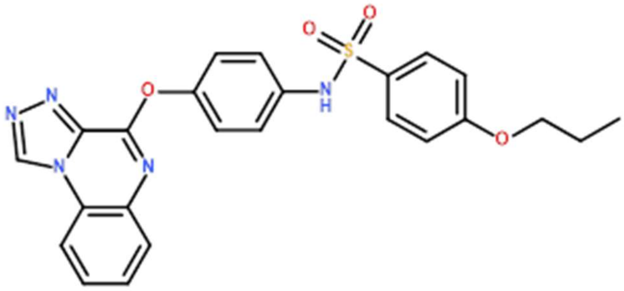 | 4.95 | 107.73 |
| 13 | 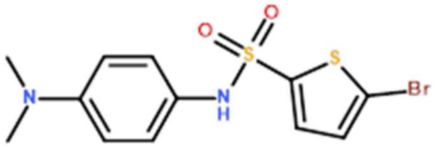 | 3.69 | 49.41 |

**B. S-Series**

| # | 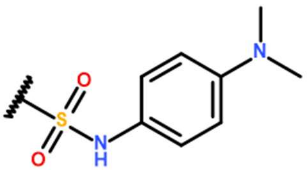 | logP | TPSA |
|---|---|---|---|
| 24 | 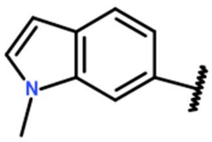 | 3.07 | 54.34 |

| | | | |
|---|---|---|---|
| **25** | 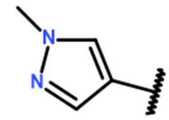 | 1.18 | 67.23 |
| **26** | 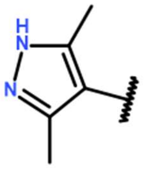 | 1.56 | 78.09 |
| **27** | 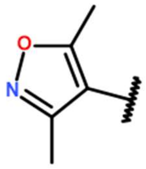 | 1.66 | 75.44 |
| **28** | 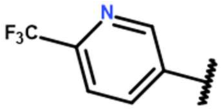 | 2.70 | 62.30 |
| **29** | 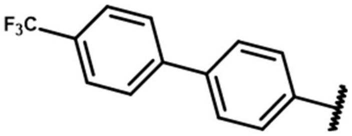 | 5.50 | 49.41 |
| **30** | 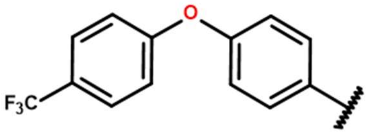 | 5.46 | 58.64 |

**C. N-series**

| **#** | 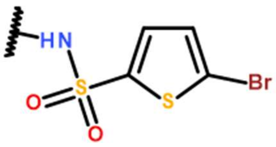 | **logP** | **TPSA** |
|---|---|---|---|
| **31** | 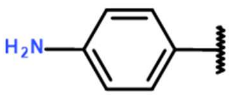 | 2.66 | 72.19 |
| **32** | 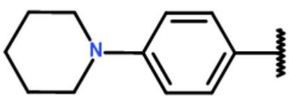 | 4.59 | 49.41 |
| **33** | 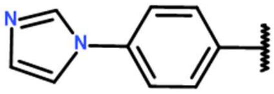 | 2.92 | 63.99 |
| **34** | 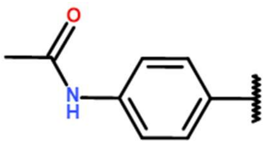 | 2.80 | 75.27 |

| | | | |
|---|---|---|---|
| **35** | 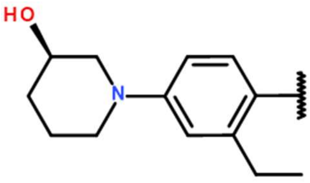 | 4.29 | 69.64 |
| **36** | 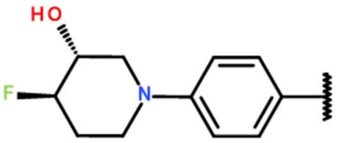 | 3.46 | 69.64 |
| **37** | 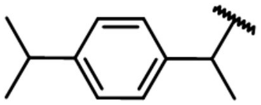 | 5.36 | 46.17 |
| **38** | 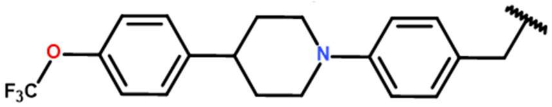 | 6.83 | 58.64 |
| **39** | 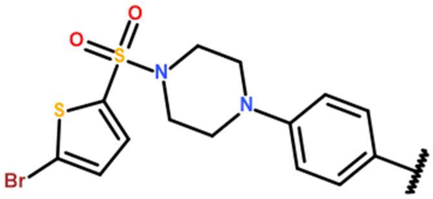 | 5.37 | 86.79 |
