## Supplemental Methods for "Identification and characterization of new structural scaffolds modulating the activity of *Mycobacterium tuberculosis* dihydroneopterin aldolase (FolB) *in vitro*"

#### Current address: R&D Center, Insol Co., Ltd., Hanam, Gyeonggi, Republic of Korea

**Keywords**

### Supplementary methods

*Chemistry*

Reagents and solvents were obtained from commercial suppliers and used without purification or drying, unless otherwise stated. $^{1}H$ NMR spectrum were recorded using a Bruker instrument at 400 MHz. TMS was used as an internal standard. LC-MS analysis were performed using an Acquity UPLC column (BEH C18, 2.1×50 mm, 1.7 μm), under the following conditions: mobile phase A: 0.05% TFA in Water; mobile phase B: 0.05% TFA in Acetonitrile; gradient (T/% B): 0/50, 3/90, 5/90, 5.01/50; flow rate: 0.4 mL/min; Temperature: 35°C. All raw NMR and HPLC data are available upon request to the corresponding author.

*Note*: numbering for hit compounds in this section are following the red labels indicated in **Figure S2**.

General procedure for the synthesis of Cluster I (compounds **Ia-c**)

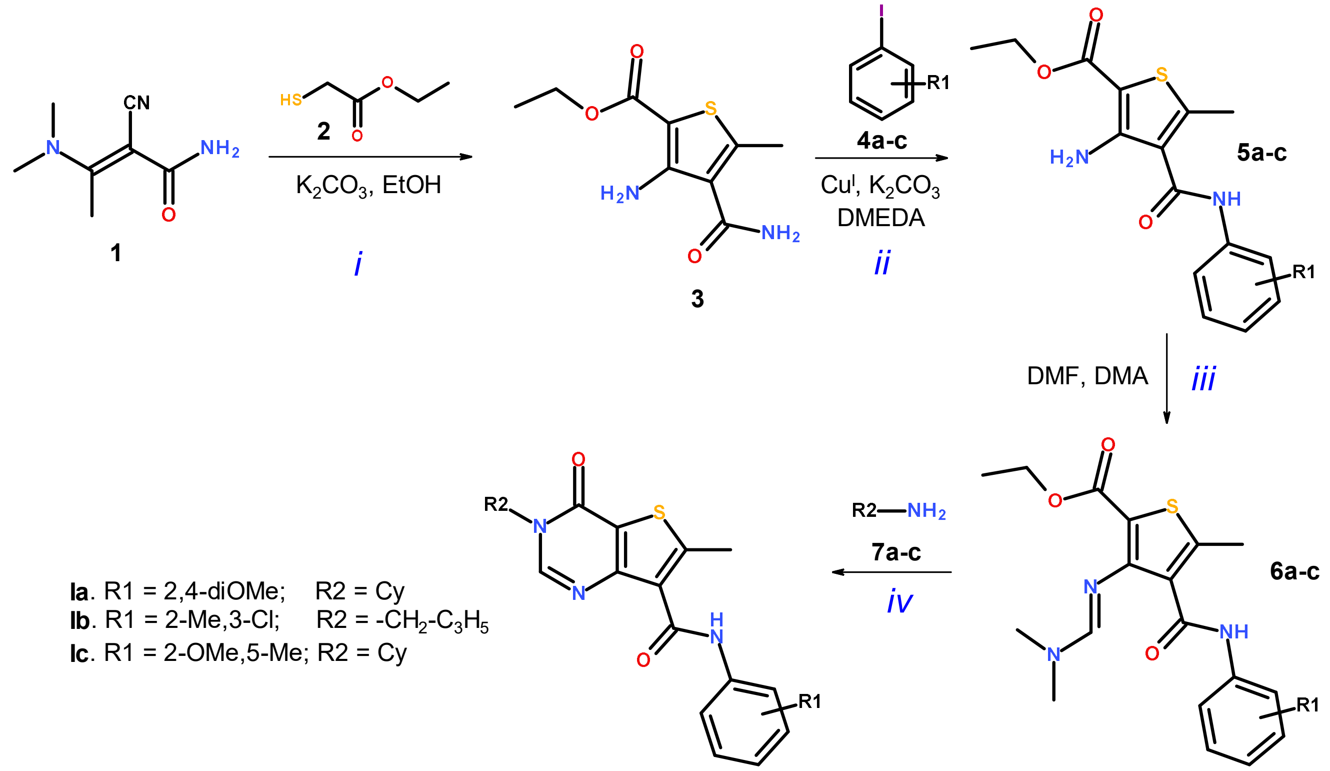

*i*. To a stirred solution of **1** (5.00 g, 33.0 mmol) and **2** (4.75 g, 39.6 mmol) in EtOH (50.0 mL) was added $K_2CO_3$ (0.318 g, 2.30 mmol). The resulting reaction mixture was stirred at 80°C for 16h. Reaction progress was monitored by TLC (40% EtOAc in petroleum ether (PE)). After completion of the reaction, the mixture was poured into ice-cold water. The precipitate formed was filtered off and washed with water and PE to afford the desired product **3** as a white solid (4.50 g, 59.8% yield).

*ii*. To a stirred solution of **3** (1 eq., ~0.5 g) in toluene (10 mL) was added CuI (0.05 eq.), $K_2CO_3$ (2.0 eq.), *N*,*N'*-dimethylethylenediamine (DMEDA, 0.10 eq.) and the corresponding iodobenzene derivative **4a-c** (0.8 eq.) at room temperature. The resulting reaction mixture was stirred at 110°C for 16h. Reaction progress was monitored by TLC (**5a**: 40% EtOAc in PE, $R_f$ = 0.6; **5b**: 30% EtOAc in PE, $R_f$ = 0.8; **5c**: 40% EtOAc in PE, $R_f$ = 0.8). After depletion of the starting material, the reaction mixture was poured into ice-cold water (50 mL) and extracted with EtOAc (2 × 100 mL). The combined organic layers were dried over anhydrous $Na_2SO_4$, filtered and concentrated under reduced pressure. The obtained crude compound was purified by silica gel column (100-200 mesh) using 15% EtOAc in PE (**5a**), 20% EtOAc in PE (**5b**) or 8% EtOAc in PE (**5c**) as eluent to afford **5a-c** as off white to pale yellow solids (48-67% yield).

*iii*. To a stirred solution of **5a-c** (1.0 eq., ~0.20 g) in ethanol (10 mL) was added *N*,*N*-dimethylformamide dimethyl acetal (DMF-DMA; 2.0 eq.) at room temperature. The resulting reaction mixture was stirred at 80°C for 4h. Reaction progress was monitored by TLC (**6a**: 20% EtOAc in PE, $R_f$ = 0.4; **6b**: 30% EtOAc in PE, $R_f$ = 0.3; **6c**: 20% EtOAc in PE, $R_f$ = 0.3). After depletion of the starting material, the reaction mixture was concentrated under reduced pressure to afford **6a-c** as pale yellow solids. Crude compounds were used in the next reaction without further purification.

*iv*. A 20 mL sealed tube was charged with **6a-c** (1 eq., ~0.150 g), the corresponding amine **7a-c** (2 eq.) and acetic acid (120 µL) at room temperature. The resulting reaction mixture was stirred at 160°C for 3h. Reaction progress was monitored by TLC (**7a**: 40% EtOAc in PE, $R_f$ = 0.4; **7b**: 30% EtOAc in PE, $R_f$ = 0.2; **7c**: 30% EtOAc in PE, $R_f$ = 0.2). After depletion of the starting material, the reaction was quenched with saturated sodium bicarbonate solution (50 mL) and extracted with EtOAc (2 × 30 mL). The combined organic layers were dried over anhydrous $Na_2SO_4$, filtered and concentrated under reduced pressure. The obtained crude compound was purified by silica gel column (100-200 mesh) using 10% EtOAc in PE (**4a,b**) or 15% EtOAc in PE (**4c**) as eluent to afford **Ia-c** as off white to pale yellow solids (27-52% yield).

3-cyclohexyl-N-(2,4-dimethoxyphenyl)-6-methyl-4-oxo-thieno[3,2-d]pyrimidine-7-carboxamide (**Ia**)

Off white solid. $^{1}$H NMR (400 MHz, DMSO-*d6*) *δ*: 1.23-1.29 (m, 1H), 1.33-1.45 (m, 2H), 1.69-1.72 (d, *J* = 12 Hz, 1H), 1.89-1.99 (m, 6H), 2.95 (s, 3H), 3.77 (s, 3H), 3.97 (s, 3H), 4.64-4.69 (m, 1H), 6.51-6.54 (dd, *J* = 2.8, 8.8 Hz, 1H), 6.68 (d, *J* = 2.4 Hz, 1H), 8.28-8.31 (dd, *J* = 1.6, 8.8 Hz, 1H), 8.76 (s, 1H), 11.58 (s, 1H). LC-MS: RT, 1.92, $[M+H]^{+}$ 428.23 m/z.

N-(3-chloro-2-methyl-phenyl)-3-(cyclopropylmethyl)-6-methyl-4-oxo-thieno[3,2-d]pyrimidine-7-carboxamide (**Ib**)

Pale yellow solid. $^{1}$H NMR (400 MHz, DMSO-*d6*) *δ*: 0.47-0.5 (m, 2H), 0.52-0.55 (m, 2H), 1.29-1.33 (m, 1H), 2.46 (s, 3H), 2.94 (s, 3H), 3.92-3.94 (d, *J* = 7.2 Hz, 2H), 7.23-7.29 (m, 2H), 8.04-8.06 (dd, J = 2.4, 6.8 Hz, 1H), 8.80 (s, 1H), 11.30 (s, 1H). LC-MS: RT 2.11, $[M+H]^{+}$ 388.13 m/z.

3-cyclohexyl-N-(2-methoxy-5-methyl-phenyl)-6-methyl-4-oxo-thieno[3,2-d]pyrimidine-7-carboxamide (**Ic**)

Off white solid. $^{1}$H NMR (400 MHz, DMSO-*d6*) *δ*: 1.23-1.33 (q, *J* = 13.1 Hz, 1H), 1.39-1.48 (q, *J* = 12.5 Hz, 2H), 1.69 (d, *J* = 12.4 Hz, 1H), 1.87-1.99 (m, 6H), 2.27 (s, 3H), 2.96 (s, 3H), 3.96 (s, 3H), 4.63-4.69 (m, 1H), 6.87-6.90 (dd, *J* = 1.6, 8.4 Hz, 1H), 6.97 (d, *J* = 8 Hz, 1H), 8.32 (m, 1H), 8.76 (s, 1H), 11.74 (s, 1H). LC-MS: RT 2.38, $[M+H]^{+}$ 412.20 m/z.

General procedure for the synthesis of Cluster III (compound **IIId** and derivatives)

| N-Series (8a + 9a-j) | | S-Series (9a + 8b-h) | |
|---|---|---|---|
| 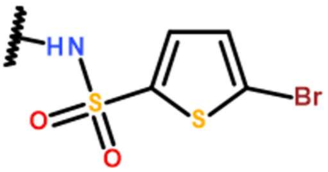 | | 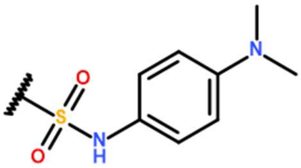 | |
| 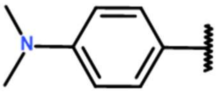 | IIId | 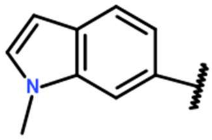 | IIIn |
| 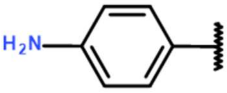 | IIIe | 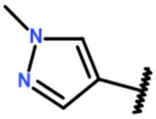 | IIIo |
| 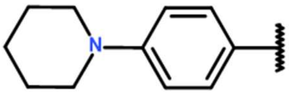 | IIIf | 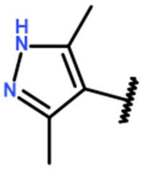 | IIIp |
| 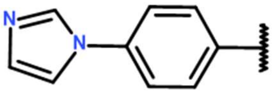 | IIIg | 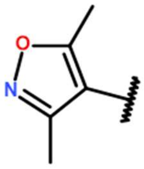 | IIIq |
| 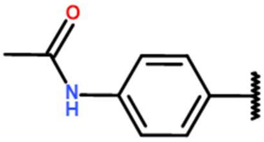 | IIIh | 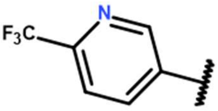 | IIIr |
| 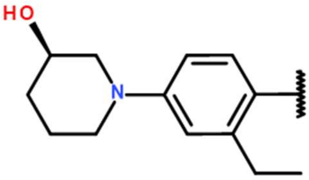 | IIIi | 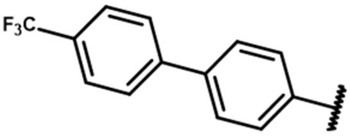 | IIIs |
| 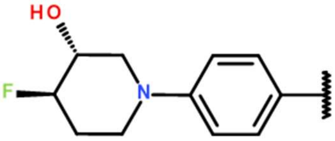 | IIIj | 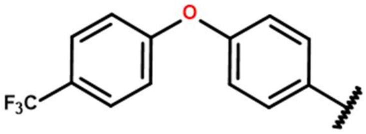 | IIIt |
| 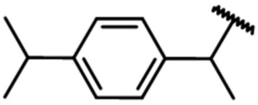 | IIIk | | |

| | |
|---|---|
| 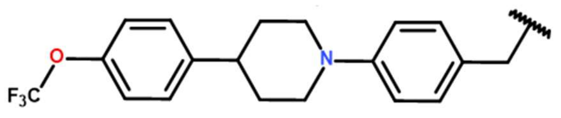 | IIIl |
| 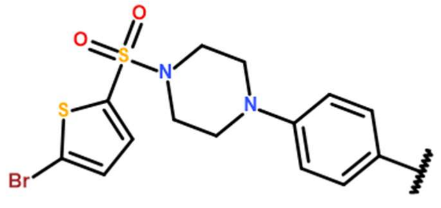 | IIIm |

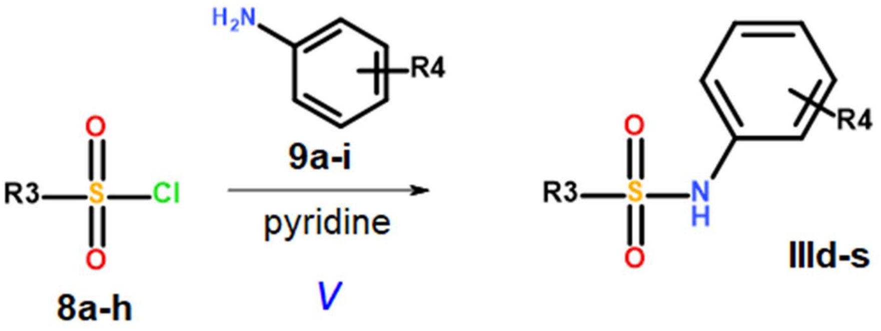

*v*. To a solution of the desired substituted heterocyclic sulfonyl chloride **8a-h** (1.2 eq.) in pyridine were added the desired heterocyclic anilines **9a-j** (1.0 eq.) and the mixture was stirred at room temperature for 2 hours. The reaction was diluted with EtOAc (20 mL), washed with water (10 mL), dried over $MgSO_4$ and concentrated under vacuum. The crude residue was purified by flash column chromatography (20% EtOAc in *n*-hexane) to afford the target compounds **IIId-m**. Compounds **IIId-m** (N-series) were obtained by reacting the same 5-bromothiophene-2-sulfonyl chloride **8a** with anilines **9a-j**. Compounds **IIIn-t** (S-series) were obtained by reacting the same N4,N4-dimethylbenzene-1,4-diamine **9a** with various sulfonyl chloride **8b-h**.

Structures of the target compounds are summarized in the table below:

<u>5-bromo-N-(4-(dimethylamino)phenyl)thiophene-2-sulfonamide</u> (**IIId**)

Brown solid, Yield 86%; $^1$H NMR (400 MHz, DMSO-*d6*) $\delta$: 2.82 (s, 6H), 7.26 (d, $J$ = 9.2 Hz, 2H), 6.93 (d, $J$ = 9.2 Hz, 1H), 7.23 (d, $J$ = 4.0 Hz, 1H), 7.60 (d, $J$ = 4.0 Hz, 1H), 9.90 (s, 1H);

MS (ESI): $[M+H]^{2+}$ 362 m/z.

N-(4-aminophenyl)-5-bromothiophene-2-sulfonamide (**IIIe**)

Brown solid; $^{1}H$ NMR (400 MHz, DMSO-*d6*) *δ*: 5.04 (s, 2H), 6.46 (d, *J* = 8.4 Hz, 2H), 6.74 (d, *J* = 8.4 Hz, 2H), 7.20 (d, *J* = 3.6 Hz, 1H), 7.27 (d, *J* = 3.2 Hz, 1H), 9.37 (s, 1H); MS (ESI): $[M+H]^{+}$ 333 m/z.

5-bromo-N-(4-(piperidin-1-yl)phenyl)thiophene-2-sulfonamide (**IIIf**)

Brown solid; Yield 82%; $^{1}H$ NMR (400 MHz, DMSO-*d6*) *δ*: 1.58-1.50 (br, 6H), 3.07 (t, *J* = 4.8 Hz, 4H), 6.85 (d, *J* = 8.8 Hz, 2H), 6.94 (d, *J* = 8.8 Hz, 2H), 7.27 (dd, *J* = 12 Hz, 8.0 Hz, 2H), 10.03 (s, 1H); MS (ESI): $[M+H]^{2+}$ 342 m/z.

N-(4-(1H-imidazol-1-yl)phenyl)-5-bromothiophene-2-sulfonamide (**IIIg**)

White solid; $^{1}H$ NMR (400 MHz, DMSO-*d6*) *δ*: 7.08 (s, 1H), 7.26 (d, *J* = 8.8 Hz, 2H), 7.31 (d, *J* = 4.0 Hz, 1H), 7.41 (d, *J* = 4.0 Hz, 1H), 7.60 (d, *J* = 8.8 Hz, 2H), 7.67 (s, 1H), 8.18 (s, 1H), 10.70 (s, 1H); MS (ESI): $[M+H]^{2+}$ 385 m/z.

N-(4-((5-bromothiophene)-2-sulfonamido)phenyl)acetamide (**IIIh**)

Brown solid; $^{1}H$ NMR (400 MHz, DMSO-*d6*) *δ*: 1.99 (s, 3H), 7.05 (d, *J* = 8.8 Hz, 2H), 7.29 (dd, *J* = 9.6 Hz, 5.6 Hz, 2H), 7.49 (d, *J* = 8.8 Hz, 2H), 9.90 (s, 1H), 10.31 (s, 1H); MS (ESI): $[M+H]^{2+}$ 376 m/z.

(R)-5-bromo-N-(2-ethyl-4-(3-hydroxypiperidin-1-yl)phenyl)thiophene-2-sulfonamide (**IIIi**)

Brown solid; $^{1}H$ NMR (400 MHz, DMSO-*d6*) *δ*: 1.28 (t, *J* = 12.4 Hz, 3H), 1.88-1.47 (m, 5H), 2.67-2.43 (m, 3H), 3.46 (d, *J* = 11.2 Hz, 1H), 3.57 (d, *J* = 9.2 Hz, 2H), 4.77 (s, 1H), 6.73-6.63 (m, 3H), 7.22 (s, 2H), 7.31 (s, 1H), 9.56 (s, 1H); MS (ESI): $[M+H]^{+}$ 445 m/z.

5-bromo-N-(4-((3R,4R)-4-fluoro-3-hydroxypiperidin-1-yl)phenyl)thiophene-2-sulfonamide (**IIIj**)

Brown solid; $^1$H NMR (400 MHz, DMSO-*d6*) δ: 1.67-1.62 (m, 1H), 2.06 (br, 1H), 2.59 (t, *J* = 12.4 Hz, 1H), 2.77 (t, *J* = 12.4 Hz, 1H), 3.57-3.52 (br, 3H), 4.45 (br, 1H), 5.36 (d, *J* = 4.4 Hz, 1H), 6.88 (d, *J* = 8.8 Hz, 2H), 6.96 (d, *J* = 8.4 Hz, 2H), 7.22 (d, *J* = 6.8 Hz, 2H), 10.06 (s, 1H); MS (ESI): $[M+H]^{2+}$ 436 m/z.

5-bromo-N-(1-(4-isopropylphenyl)ethyl)thiophene-2-sulfonamide (**IIIk**)

Brown solid; Yield 56%; $^1$H NMR (400 MHz, DMSO-*d6*) δ: 1.17(d, *J* = 6.8 Hz, 6H), 1.28 (d, *J* = 6.8 Hz, 3H), 2.83-2.79 (m, 1H), 4.40 (d, *J* = 6.8 Hz, 2H), 7.12-7.05 (m, 5H), 8.5 (s, 1H); MS (ESI): $[M+H]^{2+}$ 389 m/z.

5-bromo-N-(4-(4-(4-(trifluoromethoxy)phenyl)piperidin-1-yl)benzyl)thiophene-2-sulfonamide (**IIIl**)

Brown solid; $^1$H NMR (400 MHz, DMSO-*d6*); δ: 1.88-1.72 (m, 4H), 2.74 (t, *J* = 12.0 Hz, 3H), 3.80 (d. *J* = 12.0 Hz, 2H), 3.98 (s, 2H), 6.91 (d, *J* = 8.4 Hz, 2H), 7.10 (d, *J* = 8.4 Hz, 2H), 7.30 (d, *J* = 4.8 Hz, 3H), 7.42-7.37 (m, 3H), 8.36 (s, 1H); MS (ESI): $[M+H]^{2+}$ 576 m/z.

5-Bromo-N-(4-(4-((5-bromothiophen-2-yl)sulfonyl)piperazin-1-yl)phenyl)thiophene-2-sulfonamide (**IIIm**)

Brown solid; Yield 34%; $^1$H NMR (400 MHz, DMSO-*d6*) δ: 3.08-3.05 (br, 4H), 3.02-3.18 (br, 4H), 6.86 (d, *J* = 8.8 Hz, 2H), 6.97 (d, *J* = 8.8 Hz, 2H), 7.25 (d, *J* = 2.8 Hz, 2H), 7.48 (d, *J* = 4 Hz, 1H), 7.54 (d, *J* = 4 Hz, 1H), 10.11 (s, 1H). MS (ESI): $[M+H]^{2+}$ 627 m/z.

N-(4-(dimethylamino)phenyl)-1-methyl-1H-indole-6-sulfonamide (**IIIn**)

Brown solid; Yield 84%; $^{1}H$ NMR (400 MHz, DMSO-*d6*) $\delta$: 2.84 (s, 6H), 3.80 (s, 3H), 6.52 (d, $J$ = 8.8 Hz, 2H), 6.56 (d, $J$ = 3.2 Hz, 1H), 6.84 (d, $J$ = 8.8 Hz, 2H). 7.47-7.45 (m, 2H), 7.55 (d, $J$ = 8.8 Hz, 1H), 7.90 (s, 1H), 9.41 (s, 1H); MS (ESI): $[M+H]^+$ 330 m/z.

N-(4-(dimethylamino)phenyl)-1-methyl-1H-pyrazole-4-sulfonamide (**IIIo**)

Brown solid; $^{1}H$ NMR (400 MHz, DMSO-*d6*) $\delta$: 2.83 (s, 6H), 3.82 (s, 3H), 6.62 (d, $J$ = 8.8 Hz, 2H), 6.92 (d, $J$ = 9.2 Hz, 2H), 7.53 (s, 1H), 8.05 (s, 1H), 9.41 (s, 1H); MS (ESI): $[M+H]^+$ 281 m/z.

N-(4-(dimethylamino)phenyl)-3,5-dimethyl-1H-pyrazole-4-sulfonamide (**IIIp**)

Brown solid; Yield 38%; $^{1}H$ NMR (400 MHz, DMSO-*d6*) $\delta$: 2.09 (s, 6H), 2.82 (s, 6H), 6.60 (d, $J$ = 8.4 Hz, 2H), 6.84 (d, $J$ = 8.4 Hz, 2H), 9.18 (s, 1H), 12.72 (s, 1H); MS (ESI): $[M+H]^+$ 295 m/z.

N-(4-(dimethylamino)phenyl)-3,5-dimethyl-isoxazole-4-sulfonamide (**IIIq**)

Brown solid; $^{1}H$ NMR (400 MHz, DMSO-*d6*) $\delta$: 2.14 (s, 3H), 2.29 (s, 3H), 2.85 (s, 6H), 6.65 (d, $J$ = 8.8 Hz, 2H), 6.87 (d, $J$ = 8.8 Hz, 2H), 9.70 (s, 1H); MS (ESI): $[M+H]^+$ 296 m/z.

N-(4-(dimethylamino)phenyl)-6-(trifluoromethyl)pyridine-3-sulfonamide (**IIIr**)

Brown solid; $^{1}H$ NMR (400 MHz, DMSO-*d6*) $\delta$: 2.83 (s, 6H), 6.60 (d, $J$ = 8.8 Hz, 2H), 6.87 (d, $J$ = 8.8 Hz, 2H), 8.13 (d, $J$ = 8.4 Hz, 1H), 8.28 (d, $J$ = 8.4 Hz, 1H), 8.93 (s, 1H), 10.07 (s, 1H); MS (ESI): $[M+H]^+$ 346 m/z.

N-(4-(dimethylamino)phenyl)-4'-(trifluoromethyl)-[1,1'-biphenyl]-4-sulfonamide (**IIIs**)

Brown solid; $^1$H NMR (400 MHz, DMSO-*d6*) $\delta$: 2.80 (s, 6H), 6.59 (d, *J* = 8.8 Hz, 2H), 6.87 (d, *J* = 8.8 Hz, 2H), 7.79 (d, *J* = 8.0 Hz, 2H), 7.86 (d, *J* = 8.0 Hz, 2H), 7.95-7.90 (m, 4H), 9.74 (s, 1H); MS (ESI): $[M+H]^+$ 421 m/z.

N-(4-(dimethylamino)phenyl)-4-(4-(trifluoromethyl)phenoxy)benzenesulfonamide (**IIIt**)

Brown solid; $^1$H NMR (400 MHz, DMSO-*d6*); $\delta$: 2.81 (s, 6H), 6.60 (d, *J* = 8.8 Hz, 2H), 6.87 (d, *J* = 8.8 Hz, 2H), 7.20 (d, *J* = 8.4 Hz, 2H), 7.26 (d, *J* = 8.4 Hz, 2H), 7.70 (d, *J* = 8.4 Hz, 2H), 7.81 (d, *J* = 8.4 Hz, 2H), 9.62 (s, 1H); MS (ESI): $[M+H]^+$ 437 m/z.

General procedure for the synthesis of Cluster IV (compounds **IVa-c**)

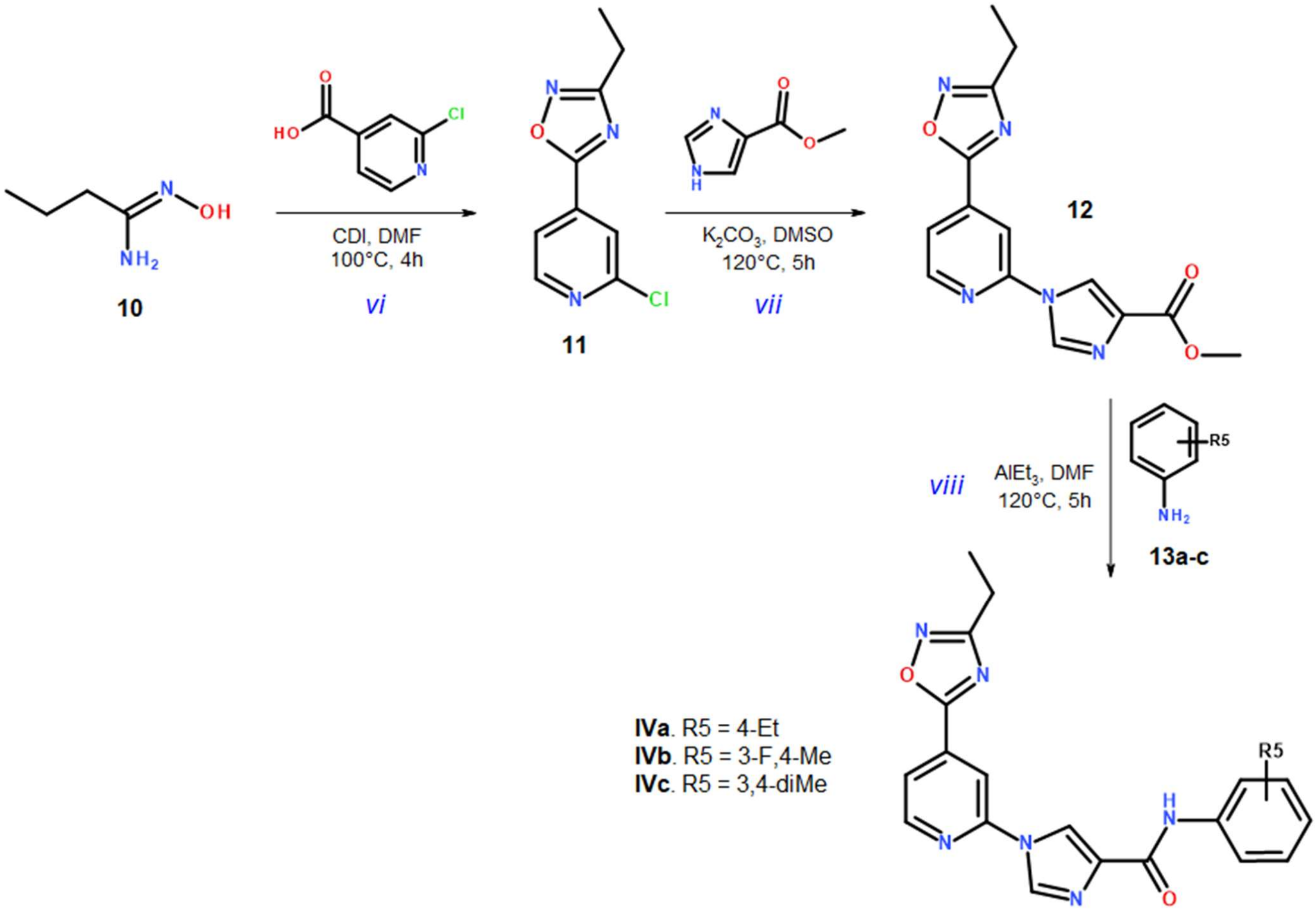

*vi*. A mixture of 2-chloroisonicotinic acid (1.0 equiv.), carbonyldiimidazole (CDI, 2.5 equiv.) and the desired amidoxime **10** (2.0 equiv.) in DMF (5 mL) was heated at 100°C for 4 hours to produce the 2-chloroisonicotinic oxadiazole **11**. After reaction completion, the mixture was cooled down to room temperature, washed with EtOAc, water and the solvent was evaporated. The crude was purified by flash column chromatography (15% EtOAc in *n*-hexane) to give compound **11**.

*vii*. A mixture of compound **11** (1.0 equiv.), $K_2CO_3$ (3.0 equiv.), methyl 4-imidazolecarboxylate (2.0 equiv.) and KI (0.1 equiv.) in DMSO (5 mL) was heated at 120°C for 5 hours to afford compound **12**. After reaction completion, the mixture was cooled down to room temperature, washed with EtOAc, water and the solvent was evaporated. Compound **12** was obtained after purification of the crude by flash column chromatography (20% EtOAc in *n*-hexane).

*viii*. A mixture of compound **12** (1.0 equiv.), triethylaluminium (2.0 equiv.) and the desired aniline **13a-c** (1.0 equiv.) in DMF (5 mL) was heated at 120°C for 5 hours. After reaction completion, the reaction mixture was cooled down to room temperature, washed with EtOAc, water and the solvent evaporated. The crude was purified by flash column chromatography (50% EtOAc in *n*-hexane) to give the target compounds **IVa-c**.

1-(4-(3-ethyl-1,2,4-oxadiazol-5-yl)pyridin-2-yl)-N-(4-ethylphenyl)-1H-imidazole-4-carboxamide (**IVa**)

White solid; $^1$H NMR (400 MHz, DMSO-*d6*) *δ*: 1.87 (t, *J* =7.6 Hz, 3H), 1.34 (t, *J* = 7.6 Hz, 3H), 2.58 (dd, *J* = 14.8 Hz, 7.6 Hz, 2H), 2.90 (dd, *J* = 14.8 Hz, 7.6 Hz, 2H), 7.18 (d, *J* = 8.4 Hz, 2H), 7.77 (d, *J* = 8.4 Hz, 2H), 8.07 (d, *J* = 4.0 Hz, 1H), 8.62 (s, 1H), 8.74 (s, 1H), 8.83 (d, *J* = 4.8 Hz, 1H), 8.89 (s, 1H), 9.95 (s, 1H); MS (ESI): $[M+H]^+$ 389 m/z.

1-(4-(3-ethyl-1,2,4-oxadiazol-5-yl)pyridin-2-yl)-N-(3-fluoro-4-methylphenyl)-1H-imidazole-4-carboxamide (**IVb**)

White solid; $^1$H NMR (400 MHz, DMSO-*d6*) *δ*: 1.34 (t, *J* = 7.2 Hz, 3H), 2.20 (s, 3H), 2.90 (dd, *J* = 14.8 Hz, *J* = 7.2 Hz, 2H), 7.22 (t, *J* = 8.4 Hz, 1H), 7.60 (d, *J* = 8.4 Hz, 1H), 7.79 (d, *J* = 12.4 Hz, 1H), 8.07 (d, *J* = 4.4 Hz, 1H), 8.62 (s, 1H), 8.76 (s, 1H), 8.83 (d, *J* = 4.4 Hz, 1H), 8.89 (s, 1H), 10.19 (s, 1H); MS (ESI): $[M+H]^+$ 393 m/z.

N-(3,4-dimethylphenyl)-1-(4-(3-ethyl-1,2,4-oxadiazol-5-yl)pyridin-2-yl)-1H-imidazole-4-carboxamide (**IVc**)

White solid, Yield 68%; $^{1}H$ NMR (400 MHz, DMSO-*d6*) *δ*: 1.34 (t, *J* = 7.6 Hz, 3H), 2.19 (s, 3H), 2.22 (s, 3H), 2.90 (dd, *J* = 14.8 Hz, 7.2 Hz, 2H), 7.09 (d, *J* = 8.4Hz, 1H), 7.57 (d, *J* = 8.4Hz, 1H), 7.64 (s, 1H), 8.06 (d, *J* = 4.8 Hz, 1H), 8.61 (s, 1H), 8.72 (s, 1H), 8.84(d, *J* = 4.8 Hz, 1H), 8.89 (s, 1H), 9.82 (s, 1H); MS (ESI): $[M+H]^+$ 389 m/z.

General procedure for the synthesis of Cluster V (compounds **Va**, **Vb**)

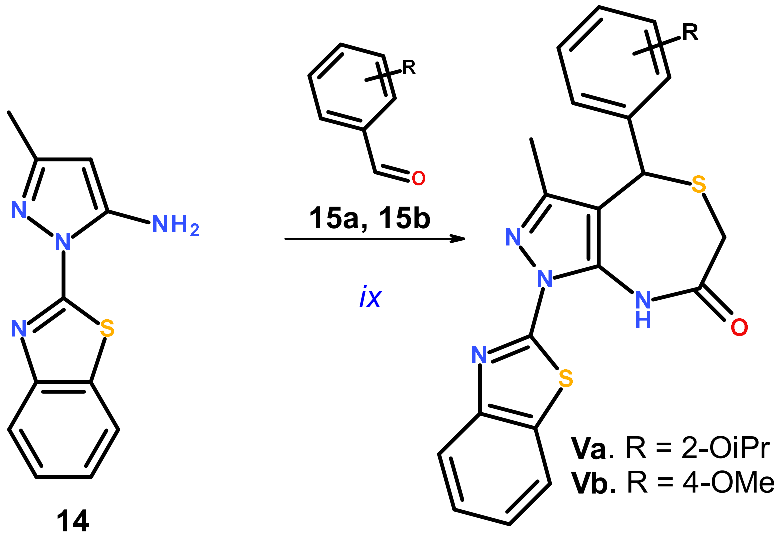

*ix*. A 10 mL microwave vial was charged with compound **14** (1 eq., ~0.50 g), **15a** or **15b** (1 eq.) and thioglycolic acid (1 eq.) at room temperature. The resulting reaction mixture was stirred at 120°C for 15-20 min under microwave irradiation. The reaction progress was monitored by TLC (20% EtOAc in PE, **Va**: $R_f$ = 0.3; **Vb**: $R_f$ = 0.4). After completion of the reaction, the crude reaction mixture was purified by silica gel column (100-200 mesh) using 10% EtOAc in PE as eluent to afford **Va** (11% yield) or **Vb** (14% yield) as off white solids.

1-(1,3-benzothiazol-2-yl)-4-(2-isopropoxyphenyl)-3-methyl-4,8-dihydropyrazolo[3,4-e][1,4]thiazepin-7-one (**Va**)

Off white solid. $^1$H NMR (400 MHz, DMSO-*d6*) $\delta$: 1.32-1.34 (d, $J$ = 4.8 Hz, 6H), 1.74 (s, 3H), 3.82-3.49 (q, $J$ = 14.1 Hz, 2H), 4.73-4.79 (m, 1H), 5.59 (s, 1H), 6.83-6.87 (t, $J$ = 7.4 Hz, 1H), 7.06-7.12 (q, $J$ = 7.9 Hz, 2H), 7.26-7.29 (t, $J$ = 7.0 Hz, 1H), 7.43-7.46 (t, $J$ = 7.6 Hz, 1H), 7.54-7.58 (t, $J$ = 7.6 Hz, 1H), 7.95 (d, J = 8.0 Hz, 1H), 8.10 (d, J = 7.6 Hz, 1H), 11.00 (s, 1H). LC-MS: RT 2.71, $[M+H]^+$ 451.44 m/z.

1-(1,3-benzothiazol-2-yl)-4-(4-methoxyphenyl)-3-methyl-4,8-dihydropyrazolo[3,4-e][1,4]thiazepin-7-one (**Vb**)

Off white solid. $^{1}H$ NMR (400 MHz, DMSO-*d6*) $\delta$: 1.82 (s, 3H), 3.26 (s, 1H), 3.47 (d, $J$ = 15.6 Hz, 1H), 3.75 (s, 3H), 5.54 (s, 1H), 6.91 (d, $J$ = 8.8 Hz, 2H), 7.24 (d, $J$ = 8.4 Hz, 2H), 7.43-7.47 (td, $J$ = 1.0, 7.7 Hz, 1H), 7.53-7.58 (td, $J$ = 1.2, 7.6 Hz, 1H), 7.95 (d, $J$ = 7.6 Hz, 1H), 8.10 (d, $J$ = 7.2 Hz, 1H), 11.03 (s, 1H). LC-MS: RT 2.70, $[M+H]^{+}$ 423.23 m/z.
